# Cell-surface proteomic profiling identifies CD72 as a regulator of microglial tiling

**DOI:** 10.1101/2025.06.02.657480

**Authors:** Tamara C. Chan, Mohit Rastogi, Micah X. Williams, Samuel Zhang, Sophia M. Shi, Steven R. Shuken, Teresa Bartling, Katleen Wild, Micaiah Atkins, Oliver Hahn, Joao A. Paulo, Saša Jereb, S. Andrew Shuster, Yongjin Yoo, Alan Napole, Victoria G. Hernandez, Liqun Luo, Marion S. Buckwalter, Beth Stevens, Benjamin E. Deverman, Deborah Kronenberg-Versteeg, Steven P. Gygi, Tony Wyss-Coray, Marius Wernig

## Abstract

Microglial tiling—the phenomenon of consistent cell-to-cell distances and non-overlapping processes—is regarded as a qualitative indicator of homeostasis, but mechanisms of microglial tiling are unknown. We used cell-surface proximity labeling and mass spectrometry to profile the microglial cell-surface proteome in an *in vitro* model of homeostatic glial physiology and used single-cell RNA sequencing and public databases to identify candidate cell-surface proteins that might modulate tiling. We designed an image-based functional assay which measures six morphological/spatial readouts to screen these proteins for modulation of tiling. CD72, a coreceptor to the B cell receptor that is expressed by microglia, disrupted tiling; we validated its effects *in vitro* and *in situ* in organotypic hippocampal brain slices. Phosphoproteomic studies revealed that CD72 modulates pathways associated with cell adhesion, repulsive receptors, microglial activation, and cytoskeletal organization. These results lay the groundwork for further investigation of the functional roles of tiling in homeostasis and disease.

## INTRODUCTION

Microglia are resident macrophages of the central nervous system (CNS) that play key roles in homeostatic upkeep of healthy brain tissue and mounting immune responses upon injury or infection^1^. Advances in single-cell RNA sequencing (scRNAseq) have revealed a vast heterogeneity of states that microglia can adopt, with many molecular signatures being specific to certain pathologies^2–4^. Not only do microglia change their transcriptional state in response to brain challenges, but also single microglial cells can adopt vastly different morphological phenotypes, including ramified, hypertrophic, and amoeboid, which are thought to be correlated to their functional state^5^. This remarkable plasticity between homeostatic and disease states, along with the fact that microglia can be replaced to restore function, has drawn great interest in recent years in therapeutically targeting microglia for brain diseases^6–8^. However, it remains imperative to understand how microglia behave in homeostatic conditions, thus informing how we can harness those processes to promote healthy microglial function.

Microglia arise from embryonic progenitors in the yolk sac that colonize the CNS, then perpetually co-exist with neuroepithelial-derived cells such as neurons, astrocytes, and oligodendrocytes^9,10^. After colonization, microglia establish a stable tiling pattern across the brain, in which cell bodies remain stationary with a consistent 50 to 60 µm distance between, yet have highly branched and motile processes that continuously extend and retract into the unoccupied space^11^. Despite this constant and seemingly random movement, microglia display territorial behavior in which processes of neighboring microglia do not overlap and are hypothesized to mutually repel^11–13^. These phenomenon—stable cell body spacing, non-overlapping processes, and interdigitated arrangement—have been collectively termed microglial tiling. Given that their homeostatic functions of scavenging debris, pruning synapses, and defending against threats all require surveillance of the tissue environment, one could reason that tiling has evolved as a means to minimize redundancy and maximize spatial coverage and surveillance rate^14^. Thus, microglial tiling is often regarded qualitatively a sign of overall microglial homeostasis.

The association between tiling and homeostasis is highlighted by observations that during disease or injury, microglial tiling is disrupted. Although microglial tiling has not explicitly been studied in any disease context, published images of brains subjected to ischemic stroke and spinal cords subjected to nerve injury display altered neighbor distances and overlaps^15,16^. This reflects known abilities of microglia to proliferate, migrate, and recruit process extensions, which presumably force microglia closer together, cause abandoned territories, and lead to overlapping processes^16–19^. We hypothesize that there are active molecular mechanisms that facilitate switching between tiling and non-tiling states. By studying how microglia constitutively regulate tiling in homeostasis, we aim to discover mechanisms that could potentially be leveraged to promote the healthy function of microglia in disease.

Here, we describe the use of an *in vitro* profiling and screening approach to address this scientific gap. We used cell-surface proteomics to profile the microglial cell-surface proteome, used RNA sequencing and publicly available databases to classify microglial cell-surface proteins, and then performed an image-based functional screen to identify cell-surface proteins (CSPs) that regulate microglial tiling.

## RESULTS

### Microglial cell-surface proteome revealed by proximity labeling and mass spectrometry

To profile the microglial cell-surface proteome, we crossed Cx3cr1-Cre^ERT2^-EYFP mice to Cre-iPEEL mice to generate mice with microglia-specific, plasma membrane-targeted, hemagglutinin (HA)-tagged horseradish peroxidase (HRP) expression^20,21^. On performing proximity labeling in acute brain sections, we observed no biotinylation on microglia, despite positive HRP expression as visualized by immunostaining for HA-tag (Figure S1A). However, microglial cell-surface labeling did succeed after isolating mixed glia from early postnatal brains of the same mice, a previously uncharacterized *in vitro* model in which tiling is preserved (Figure 1A). Thus, we induced microglia-specific HRP expression *in vitro* with 4-OHT and performed cell-surface proximity labeling (Figure 1B). Immunostaining for HA-tag showed specific expression only within EYFP-positive cells and staining for biotinylated protein revealed specific biotinylation only on HRP-expressing cells (Figure 1C). Furthermore, we restricted immunodetection to the cell surface by staining without permeabilization, thus confirming cell-surface labeling (Figure 1D). As expected, biotinylation was not observed in control cultures untreated with H_2_O_2_ (+HRP/-H_2_O_2_ Control 1) (Figure 1E). Furthermore, we did not detect EYFP, HA, or biotinylated signal in control cultures derived from Cx3cr1-Cre^ERT2^-EYFP -/-; Cre-iPEEL littermate controls (-HRP/+H_2_O_2_ Control 2) (Figure 1E). Next, we prepared lysates from the experimental and two negative control conditions, performed streptavidin bead enrichment, and performed liquid chromatography-tandem mass spectrometry (LC-MS/MS) to identify and quantify enriched CSPs (Figures 1B and 1E-F). In total, we detected 2064 proteins (Figure 1F). To identify CSPs, we employed a previously published ratiometric cutoff strategy pairing the experimental condition with each of the negative control conditions, which distinguishes enriched proteins from non-biotinylated background^21^ (Figure 1F). In a receiver operating characteristic analysis, the top 25% most highly enriched proteins yielded nearly vertical curves, confirming highly specific enrichment in each pairing (Figure S1B). Gene ontology (GO) cellular component enrichment analysis of the top 150 proteins with highest ratios yielded terms such as plasma membrane, cell surface, and intrinsic component of plasma membrane, confirming the enrichment of the expected cell compartment (Figure 1G). Ratio cutoffs were determined as where the true-positive rate minus false-positive rate was maximized^21^ (Figure S1C, STAR Methods). The post-cutoff, intersected, filtered list resulted in 241 CSPs, which we call the microglial cell-surface proteome (Figure 1F, Table S1). GO biological process enrichment analysis of 241 CSPs revealed enriched terms such as cell adhesion, multicellular organism development, locomotion, and cell differentiation—highlighting the importance of the cell surface in cell-to-cell and cell-to-environment communication (Figure 1H). Notably, one enriched GO term was neurogenesis, which we found to include known axon guidance molecules, suggesting that microglia could potentially use axon guidance mechanisms to sense neighboring microglia and repel each other, thus maintaining separate territories (Figure 1H).

**Figure 1.**
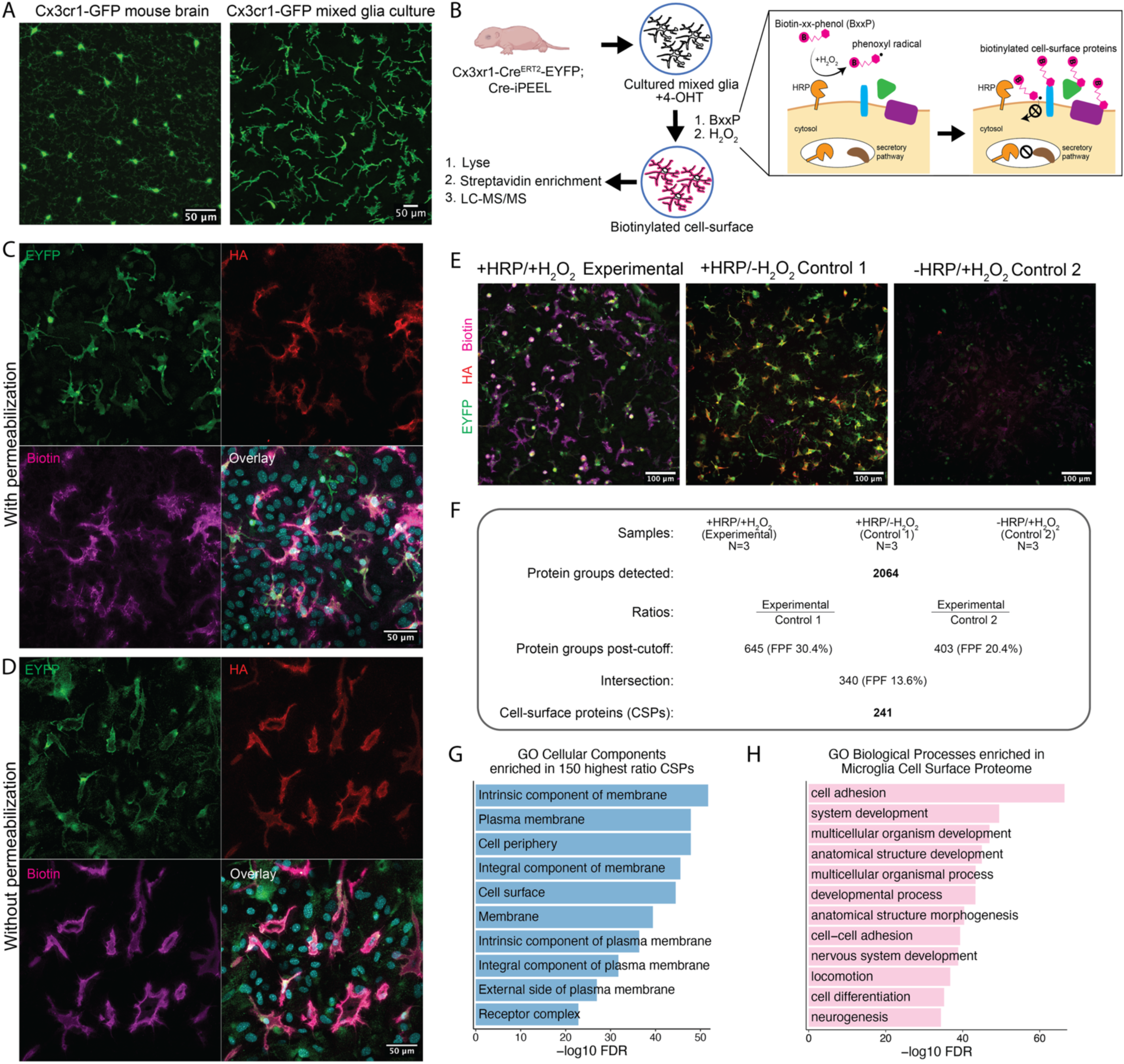
Microglial cell-surface proteome revealed by proximity labeling and mass spectrometry. **(A)** Immunofluorescent images of microglial tiling in mouse brain section (left) and mixed glia culture (right), from Cx3cr1-GFP mice, stained for GFP. **(B)** Workflow for proximity labeling of mixed glia culture, including schematic of HRP-mediated catalysis of cell-surface biotinylation. **(C and D)** Immunofluorescent images showing co-localization of microglia (stained for EYFP, green), HRP expression (stained for HA-tag, red), and biotinylated protein (stained with Streptavidin-647, magenta); stained with 0.2% Triton-X in all buffers (C) or without Triton-X (D). **(E)** Immunofluorescent images overlaying all three channels (EYFP in green, HA in red, biotinylated protein in magenta), showing conditions: Cx3cr1-Cre^ERT2^-EYFP+ glia treated with all labeling reagents (Experimental), Cx3cr1-Cre^ERT2^-EYFP+ glia treated with labeling reagents except H_2_O_2_ (Control 1), Cx3cr1-Cre^ERT2^-EYFP-glia treated with all labeling reagents (Control 2). **(F)** Summary of ratiometric and cutoff analysis of microglial cell-surface proteome. N=3 15cm dishes for each condition were used as starting material for streptavidin enrichment and label-free LC-MS/MS. See also Figure S1B and C, Table S1, and STAR Methods. **(G)** Enriched GO Cellular Components from the top 150 proteins with highest ratios within the ranked Experimental/Control 2 list. Enrichment results were similar for the Experimental/Control 1 list (data not shown). See Table S1. **(H)** Enriched GO Biological Processes from final 241 CSPs.

### Single-cell RNA sequencing characterizes mixed glia culture and reveals cell type-specific origins of the microglial surface proteome

HRP-mediated proximity labeling is expected to label not only CSPs produced by microglia, but also proteins from cell types that physically contact microglia or secrete proteins over a longer distance that bind to microglia through receptor-ligand interactions^22^. To determine microglia-specific transcript expression of our CSPs, we performed scRNAseq on the mixed glia culture. Unsupervised clustering delineated eight clusters, in which we used differentially expressed genes (DEGs) to distinguish microglia, including microglial marker genes *Fcrls, Tmem119, P2ry12,* and *Aif1*^3^ (Figures 2A-C, Table S2). A cluster expressing those microglial markers, in addition to cell cycle markers *Top2a, Cdk1*, and *Ube2c,* was identified to be proliferative microglia^23^ (Figure 2B, Table S2). Two major astrocyte clusters—Astrocytes 1 and Astrocytes 2—expressed markers such as *Gfap, Gja1, Aldoc,* and *Aqp4*^24,25^(Figures 2B and 2D, Table S2). Oligodendrocyte progenitor cells (OPCs) were defined by markers *Pdgfra, Olig1*, and *C1ql*^26,27^ (Figure 2B, Table S2). Ependymal cells expressed canonical markers such as *Myb, Pifo,* and *Foxj1*^28^ (Figure 2B, Table S2). Furthermore, there were two less distinct clusters that shared DEGs with neuroepithelial lineage cell types but did not have canonical markers for mature brain cell types. We deemed them to likely be radial glia, identified by markers *Shh, Ntn1, and Mia*, and neuroblasts, identified by markers *Hmmr, Aspm, Pbk,* and *Top2a*^29^ (Figure 2B, Table S2). Notably, non-microglial populations had a gradient of gene expression, reflecting their immature and differentiating states at the neonatal timepoint at which the cultures were isolated.

**Figure 2.**
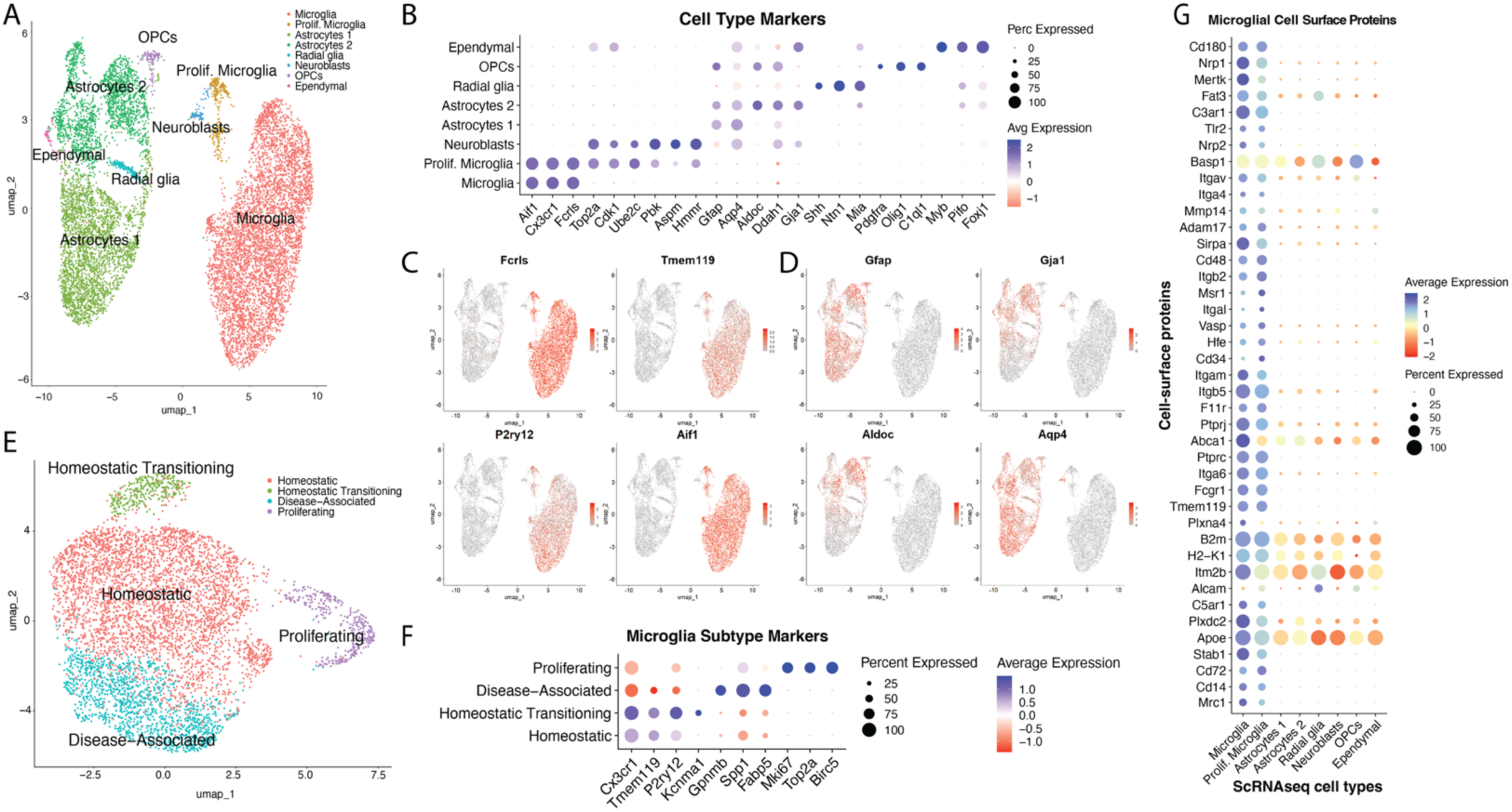
scRNAseq characterizes mixed glia culture and reveals cell type-specific origins of the microglial cell-surface proteome. **(A)** UMAP plot of all cells collected from the mixed glia culture, with clusters annotated as cell types. **(B)** Dot plot of DEG markers (each cluster over all other cells) used to assign cell types in (A). See also Table S2. **(C)** UMAP clusters overlaid with expression patterns for microglial genes. **(D)** UMAP clusters overlaid with expression patterns for astrocytic genes. **(E)** UMAP plot showing subsetted microglia cluster from (A), subclustered to reveal four microglial subtypes. **(F)** Dot plot of DEG markers (each subcluster over all other microglia) used to assign microglial subtype in (E). See also Table S3. **(G)** Dot plot of a subset of CSPs identified in Figure 1 (y-axis) whose corresponding transcripts are primarily produced by microglia based on scRNAseq expression (clusters along x-axis). See also Figure 1 and S2A, and Table S4.

Subsetting out the microglia and performing unsupervised clustering again revealed four subclusters (Figures 2E-F, Table S3). The largest cluster, comprising 64.2% of all microglia, had higher expression of classical homeostatic markers *Cx3cr1, Tmem119, P2ry12* and was labeled as Homeostatic^3^ (Figures 2E-F, Table S3). Notably, other *ex vivo* microglia culture systems exhibit downregulation/absence of these signature genes, suggesting that our mixed glia culture is more relevant to homoestatic physiology^30–32^. This agrees with their ramified morphology and tiled arrangement, which is observed *in vivo* but generally not *in vitro*^30–32^ (Figure 1A). Another microglial cluster (5.5%) appeared very similar to the Homeostatic cluster, however with the strongest differentially upregulated gene being *Kcnma1*, a ubiquitous alpha subunit of Ca^2+^-activated K^+^ (BK) channels, suggesting that these cells might be undergoing a transition or are primed towards a more activated and migratory state but have not yet downregulated homeostatic markers^33,34^. We called them “Homeostatic Transitioning” (Figure 2F, Table S3). 24.2% of microglia differentially expressed genes such as *Gpnmb, Spp1*, and *Fabp5,* which are associated with neurodegenerative diseases, thus we called this population Disease-Associated^35,36^ (Figure 2F, Table S3). Lastly, proliferating microglia (6.1%) again clustered away from the rest, with differential expression of *Mki67*, *Top2a*, and *Birc5*^23,37,38^ (Figure 2F, Table S3). Altogether, deeper characterization revealed several subtypes of microglia that were mostly homeostatic, suggesting that this mixed glia culture is appropriate for studying homeostatic processes such as tiling.

Cross-referencing to our microglial cell-surface proteomics data (Figure 1), we were able to detect 235 out of the 241 CSPs in the scRNAseq data. The most specific CSPs belonged to the microglia cluster, including Cd14 and Itgam (Cd11b), well known monocyte/macrophage markers that we would not expect to be transcriptionally expressed in any neuroepithelial lineage cells; thus, we can confidently say that these CSPs were produced by microglia^39^ (Figure 2G). However, the majority of the 241 CSPs were produced by non-microglial cells, suggesting that the microglial cell surface interacts with secreted or cell-surface proteins from all other cell types in the culture (Figure S2A, Table S4). In summary, characterization of the mixed glia culture via scRNAseq complemented the cell-surface proteomics data by allowing cell type assignment of CSPs, thus guiding our exploration of potential regulators of microglial tiling.

### An image-based screen identifies CD72 as a strong regulator of tiling

To identify molecular mechanisms of microglial tiling which could be regulated by cell-surface interactions, we designed a targeted screen to identify CSPs that play a functional role in tiling. First, we used StringDB functional annotations for all 241 CSPs to filter for those related to guidance, attraction, and repulsion, which left us with 57 proteins (Figure 3A, Table S5). Next, we sought proteins whose RNA expression changes in a condition in which tiling is disrupted: stroke^15^. Cross-referencing our 57 proteins with bulk RNA sequencing data of significantly up or downregulated microglial transcripts three days after distal middle cerebral artery occlusion stroke in mice, we were left with 26 proteins^40^ (Figures 3A-B, Table S5). We added to our list known ligands to CSP candidates, then selected several control proteins/small molecules to serve as morphological readout controls, totaling to 32 molecules (Figure 3A, Table S5). To perform the screen, we isolated mixed glia from Cx3cr1-GFP neonatal mice and exposed them to recombinant versions of our candidates, either in soluble form or embedded in Matrigel, which can recapitulate interactions that rely on extracellular matrix (ECM)^41^.

**Figure 3.**
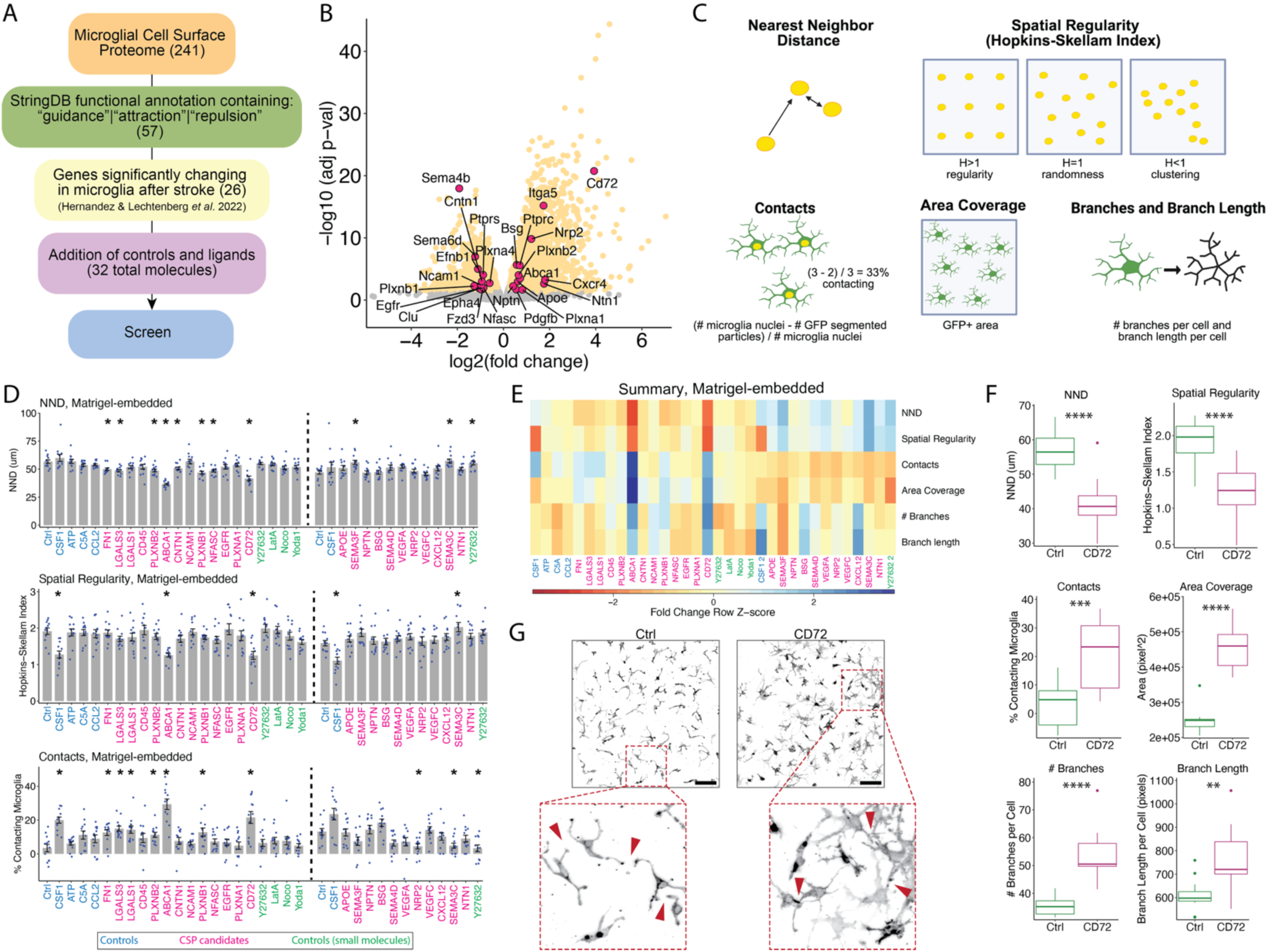
An image-based screen identifies CD72 as a strong regulator of tiling. **(A)** Tiling candidate list development. 241 CSPs were filtered by 1) StringDB functional annotations matching key words, 2) published bulk RNA sequencing dataset of significantly up- or downregulated (p-adjusted <0.05) microglial transcripts three days after distal middle cerebral artery occlusion stroke in mice. Known ligands to CSPs and controls for morphometric readouts were added. See Table S5. **(B)** Volcano plot of microglial transcripts three days after distal middle cerebral artery occlusion stroke in mice, with 26 significantly changing genes of corresponding CSPs (p-adjusted <0.05) highlighted in magenta dots. **(C)** Visual explanation of the six features quantified as the screen readout. Samples were stained for GFP (from Cx3cr1-GFP) to label microglia and DAPI. Cartoon yellow nuclei indicate that the overlap between GFP and DAPI signal was used for that quantification. See STAR Methods. **(D)** Data from NND, spatial regularity, and contacts of the Matrigel-embedded screen. N= Four images from each of three wells, 12 technical replicates. Bars show mean ± SEM. Two rounds of screening were performed, with rounds separated by vertical dotted lines. Molecules are shown in the order they were tested. X-axis text color represents category of molecule. Adjusted p-values (Benjamini-Hochberg (BH) method): *p<0.05 by Welch’s t-test between respective molecule and Ctrl (vehicle 0.1% BSA) within round. See also Figure S3A and B. **(E)** Summary heatmap of all features for every candidate in both rounds of the Matrigel-embedded screen. Color bar represents fold-change over respective Ctrl within round. X-axis text color represents the same as in (D). **(F)** Boxplots of six features showing differences between Ctrl and CD72 samples (data extracted from Matrigel-embedded screen). P values: *p<0.05, **<p<0.01, ***p<0.001, ****p<0.0001 by Welch’s t-test. **(G)** Representative images (GFP channel) for Ctrl and CD72 (Matrigel-embedded screen). Red arrowheads indicate locations of upheld tiling in Ctrl and disrupted tiling in CD72 condition. Scale bar = 100µm.

After 48 hours, cultures were fixed and stained for GFP and DAPI and we performed automated confocal imaging. Because tiling is a complex morphological process with no single characteristic molecular readout, we applied custom image analysis pipelines to measure six visual features that, collectively, could indicate a change in tiling: nearest neighbor distance (NND), spatial regularity, area coverage, contacts, number of branches per cell, and branch length per cell (Figure 3C, STAR Methods). As a positive control for morphological readouts, CSF1, a known microglial survival and proliferative cytokine, induced a change in morphology that manifested in significant changes to contacts, spatial regularity, and area coverage^42^ (Figures 3D and S3A). Another control, CCL2, a known chemotactic cytokine for microglia, did not have a large effect when embedded in Matrigel, but caused significant changes to NND, contacts, and area coverage when treated in solution, highlighting the importance of performing both versions of the screen^43^ (Figures 3D and S3B).

Amongst our candidate proteins, CD72 significantly decreased NND, decreased spatial regularity, increased contacts, increased area coverage, and increased branch number and length (Figures 3E-G). These observations indicate an invasion of neighboring territory through clustering, overlapping processes, and extending more/longer branches, together indicating a strong disruption of tiling.

### Recombinant CD72-mediated disruption of tiling is dose-dependent, reversible, and consistent in situ

We found CD72—a 45kDa type II transmembrane glycoprotein and coreceptor that negatively regulates B cell receptor (BCR) signaling^44^— particularly interesting not only due to its receptor capabilities, but also due to its known and predicted functional partners (SEMA4D and PLXNB1) that are involved in repulsive axon guidance^45–47^. In our culture, it is specifically expressed by microglia, however its function in microglia is not well studied (Figure 2G). Thus, we moved forward with CD72 as our primary candidate.

To further characterize the phenotype, we treated Cx3cr1-GFP mixed glia with a range of doses of recombinant CD72 (rCD72) from 400 ng/ml (same dose treated in screen) to 0 ng/ml (Figure 4A). Microglial tiling broadly showed clear dose-dependence, with the most disruption at the two highest doses, 400 and 40 ng/ml: microglia had closer neighbors, more clustered spatial distribution, more contacts between neighbors, covered more area, and had greater process complexity (Figure 4B). We observed a biphasic response in contacts and area coverage, with significantly reduced contacts and reduced area coverage at the two intermediate doses (4 and 0.4 ng/ml) compared to high or low doses (Figure 4B). This suggests that these two features of tiling may be oppositely regulated by exposure to intermediate or high amounts of rCD72.

**Figure 4.**
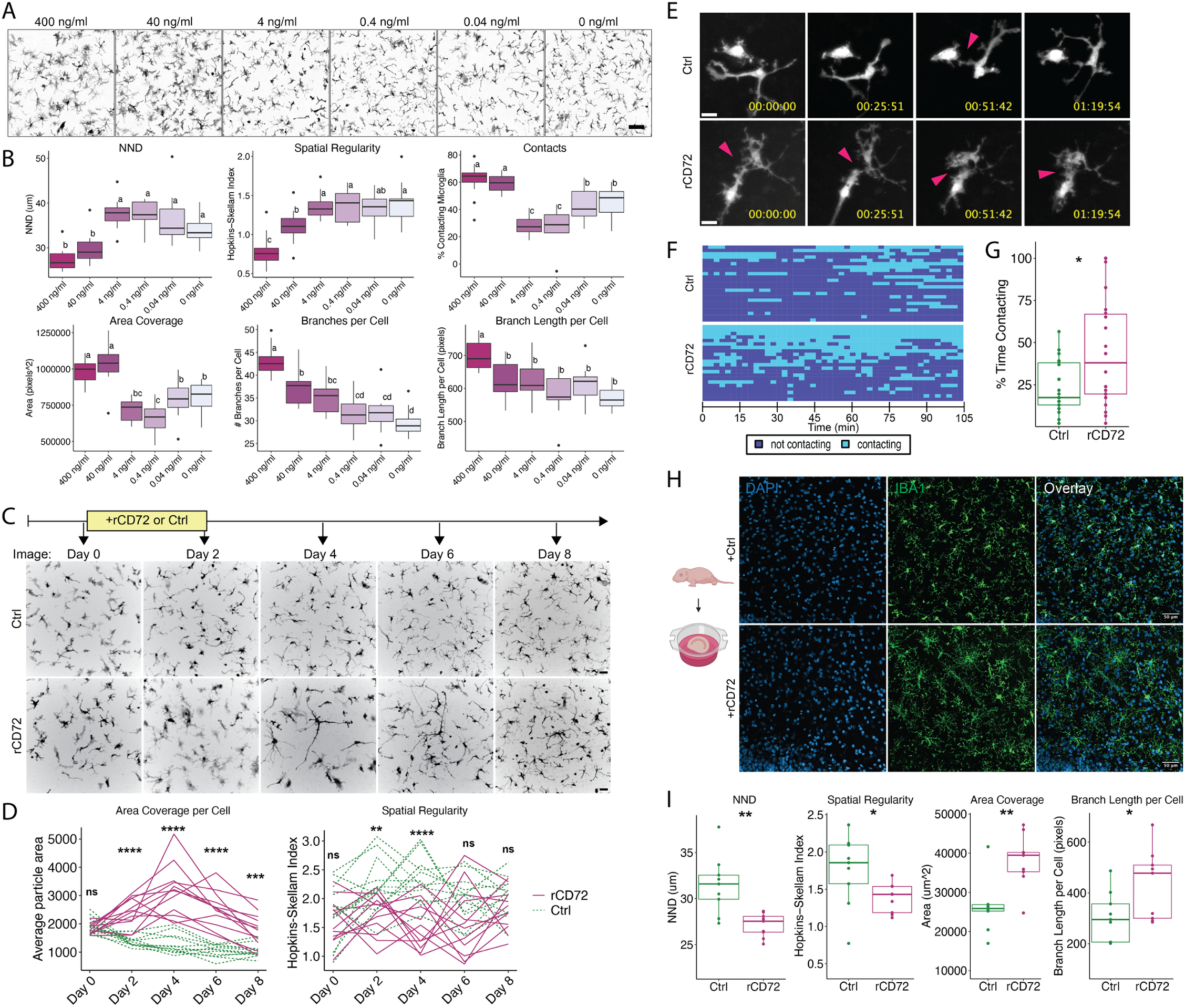
Recombinant CD72-mediated disruption of tiling is dose-dependent, reversible, and consistent *in situ*. **(A)** Representative images of GFP channel (from Cx3cr1-GFP mixed glia) treated with varying doses of rCD72. Scale bar = 100µm. **(B)** Boxplot quantifications of all features at every dose tested. N=four images in each of three wells, 12 technical replicates. One-way ANOVA, post-hoc Tukey’s test, and compact letter display was applied. **(C)** Experimental timeline and representative images from longitudinal live imaging of endogenous GFP signal before incubation (Day 0), after two days of incubation with (Day 2), and after removing (Days 4, 6, and 8) 400ng/ml soluble rCD72 or Ctrl (vehicle 0.1% BSA). Scale bar = 50µm. **(D)** Quantifications of area coverage per cell and spatial regularity at every timepoint. N=four fields of view in each of three wells, 12 technical replicates. The same fields of view were imaged across days and are connected by lines. P-values: ns = not significant, *p<0.05, **p<0.01, ***p<0.001, ****p<0.0001 by Welch’s t-test between means of Ctrl and CD72 at that timepoint. **(E)** Confocal timelapse imaging of Ctrl- and rCD72-treated (400ng/ml) Cx3cr1-GFP mixed glia over 1.75 hours (time is displayed as HH:MM:SS format). Representative images are zoomed-in fields of view imaged across time (left to right). Magenta arrowheads indicate where a contact was detected. Scale bar = 25µm. **(F)** Binary heatmap showing Ctrl, N=23, and rCD72, N=22, fields of view (each row = one field of view) across two videos per condition. Time is represented from left to right. Instances of contact, light blue; non-contact, dark blue. **(G)** Boxplot showing percent time contacting. P-value: *p<0.05 by Wilcoxon Rank Sum Test. **(H)** *In situ* treatment of wildtype organotypic hippocampal slices with Ctrl or rCD72, with representative confocal images stained for DAPI in blue and Iba1 in green. Scale bar = 50µm. **(I)** Boxplot quantifications of tiling features from images in (H), quantified from the Iba1 channel. Data represent N=9 images across three biological replicates. P-values: *p<0.05, **p<0.01 by Welch’s t-test.

Next, we asked whether the rCD72-mediated disruption in tiling is reversible. We performed longitudinal confocal live imaging before incubation (Day 0), after two days of incubation with (Day 2), and after removing rCD72 (Days 4, 6, and 8) (Figure 4C). We then quantified the area coverage per cell and spatial regularity using only the endogenous GFP signal. rCD72 treatment caused cells to become significantly enlarged and irregularly spaced after two days of incubation relative to control wells (Figure 4D). Even larger effects were observed at Day 4 (although the protein was absent between Days 2 and 4), suggesting that rCD72 causes a lasting response and that peak tiling disruption occurs four days after initial exposure (Figure 4D). Both features began converging with control at Day 6 and more so at Day 8, when area coverage per cell remained significantly different from control but to a lesser degree than Days 4 and 6 and spatial regularity was not different from control (Figure 4D). Thus, translocation of cells to restore spatial regularity happens on a faster time scale than reversal of cell area, and these features are likely regulated through different pathways in response to rCD72.

Non-overlapping processes is one of the facets of microglial tiling *in vivo*, however, in the mixed glia culture, we do observe contacts even in control conditions; these are increased upon exposure to rCD72 (Figure 3F). To ask if lack of repulsion drives this phenotype, we performed live confocal timelapse imaging and looked specifically for instances of cell-cell contact within both control and rCD72 conditions. In control conditions, contacts between processes of neighboring cells mostly occurred briefly, suggesting contact-dependent repulsion (Figure 4E). In the rCD72 condition, we observed that instances of contact were more persistent (Figures 4E-F). The average amount of time spent contacting was increased in the rCD72 condition, with some contact locations lasting 100% of the time (Figure 4F-G). Altogether, these data suggest that rCD72 treatment attenuates contact-dependent repulsion of neighboring microglial processes.

To confirm our results in a more physiologically relevant setting, we prepared organotypic hippocampal brain slices from wildtype neonatal mice and cultured them in the presence of rCD72 or control for four days (Figure 4H). We observed substantial changes in microglial tiling including significantly decreased NND, decreased spatial regularity, increased area coverage, and increased branch length (Figure 4I). These phenotypes from *in situ* hippocampal slices were consistent with our observations from *in vitro* mixed glia.

### rCD72 induces molecular pathways consistent with tiling disruption and microglial immune response

To reveal the molecular processes underlying rCD72-mediated disruption of microglial tiling, we performed LC-MS/MS on the proteome and phosphoproteome of sorted microglia from the mixed glia culture treated with rCD72 or control for 48 hours, multiplexing samples with tandem mass tags (TMT). Altogether, we quantified 7,158 proteins and 6,797 phosphorylation sites on 2,686 proteins. We identified 353 significantly upregulated proteins and 293 significantly downregulated proteins (fold change > 1.5 & adjusted p-value < 0.05) with rCD72 treatment (Figure 5A, Table S6). Within the phosphoproteomic data, we identified 376 significantly upregulated phosphorylation events on 286 proteins, and 493 significantly downregulated phosphorylation events on 350 proteins (fold change > 1.3 & adjusted p-value < 0.05) (Figure S4A, Table S6). GO enrichment analysis from the protein abundance dataset revealed a range of immune response, cell-cell communication, and cytoskeletal changes (Figure 5B).

**Figure 5.**
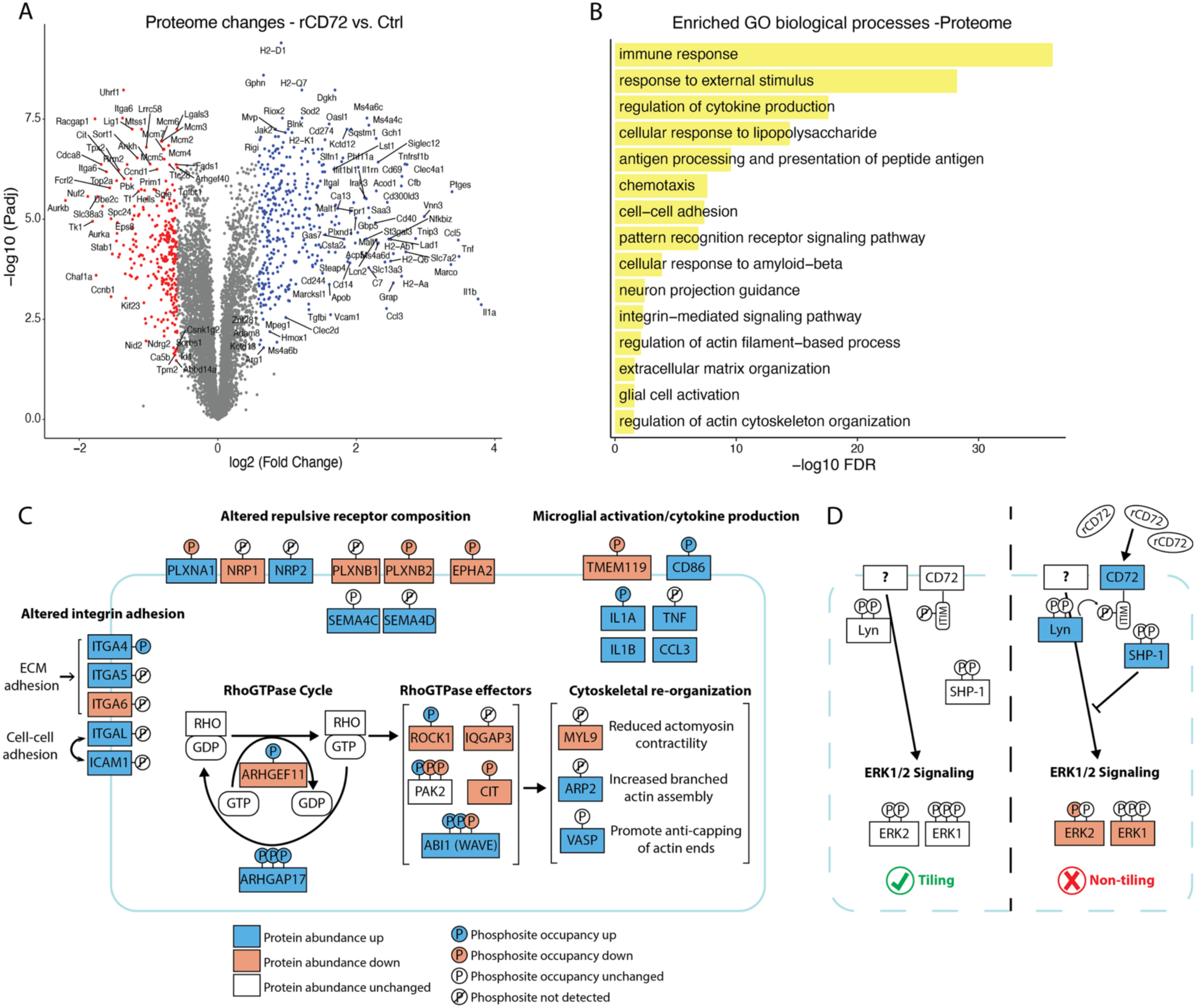
rCD72 induces molecular pathways consistent with tiling disruption and microglial immune response. **(A)** Volcano plot showing protein abundance changes in microglia enriched from the mixed glia culture treated with Ctrl (vehicle 0.1% BSA) or 400ng/ml rCD72, N=6 each. Welch’s t-test was applied between Ctrl and rCD72, then p-values adjusted using BH method. Average fold-changes were calculated. Red dots denote downregulation and blue dots denote upregulation with rCD72 treatment (fold-change cutoff:1.5, adjusted p-value cutoff: 0.05). See also Table S6. **(B)** Enriched biological processes from significantly changing proteins. **(C)** Schematic depicting specific proteins related to enriched GO terms in (B) and their phosphorylation sites. The criterion for coloring was adjusted p-value <0.05 (no fold-change cutoff). The number of phosphorylation sites depicted on each protein is not necessarily the number of phosphorylation sites detected; however, any significant changes are shown. See Table S6. **(D)** Schematic depicting proteins within known CD72 signaling pathways and hypotheses about their roles in microglial tiling. Left of the black dotted line represents proteins at steady-state: CD72 is a co-receptor to an unknown microglial receptor that is analogous to the BCR in B cells. Right of the black dotted line: upon rCD72 treatment, LYN (kinase associated with the unknown receptor) phosphorylates the ITIM motif on CD72, recruiting SHP-1 (phosphatase) to downmodulate signaling by the unknown receptor, which could result in reduced ERK1/2 signaling, resulting in non-tiling. Coloring of proteins and phosphorylation sites on the right of the dotted line follows the same criterion as in (C).

Many of the most significantly enriched GO terms were immune-related: for example, regulation of cytokine production, antigen processing and presentation of antigen, and response to lipopolysaccharide (Figure 5B). We observed downregulation of homeostatic protein TMEM119, upregulation of pro-inflammatory macrophage marker CD86, and increased production of inflammatory cytokines IL1A, IL1B, TNF, and CCL3, demonstrating activation of immune response pathways in response to rCD72 exposure^2,3,48,49^ (Figure 5C). These results suggest that the disruption of tiling may cause, require, or simply coincide with microglial immune activation.

The GO term “neuron projection guidance” was enriched due to dysregulation of axon guidance molecules including plexin, neuropilin, and ephrin receptors, as well as semaphorin ligands (Figure 5B-C). For example, PLXNB1, which was downregulated after rCD72 treatment, mediates growth cone collapse and neurite retraction upon binding its ligand SEMA4D (upregulated after rCD72 treatment)^50^. Additionally, PLXNB1 has been shown to regulate spacing of astrocytes and microglia around amyloid plaques in a mouse model of AD, with PLXNB1 knockout showing more dense and compact glial nets around the plaque^51^. Downregulation of PLXNB1, amongst other changes to the repulsive receptor landscape in response to rCD72 could be responsible for attenuating contact-dependent repulsion between microglia.

Axon guidance molecules facilitate their cellular outcomes via RhoGTPase regulation of the cytoskeleton^52^. We observed that cytoskeletal organization is an enriched pathway in both the proteome and phosphoproteome datasets (Figures 5B and S4B). Changes to RhoGTPase cycle regulators (ARHGEF11, ARHGAP17) and effectors (ROCK1, ABI1, etc.) are consistent with reduced Rho GTPase signaling via favoring the inactive, GDP-bound state^53^ (Figure 5C). For example, there was a decrease in phosphorylation at site S216 on ABI1; phosphorylation at this site has been shown to attenuate F actin assembly^54,55^ (Figure 5C). In this context, promotion of F actin assembly via de-phosphorylation of ABI1 would lead to increased actin-binding ARP2, which is indeed upregulated^54–56^ (Figure 5C). Downregulation of regulatory light chain of myosin II (MYL9) suggests reduced actomyosin contractility and subsequent cell relaxation^57^ (Figure 5C). Upregulated VASP suggests protection of growing actin filaments ends against capping, thus elongation of actin filaments^58^ (Figure 5C).

Another enriched term was integrin-mediated signaling pathway (Figure 5B). Adhesion molecules have been studied in tiling of other systems, such as DSCAM1 in *Drosophila* neurons and hepaCAM in mouse astrocytes^59,60^. We found ECM-adhesion integrins ITGA4, ITGA5, ITGA6 were changed, which could facilitate the rearrangement of cells into a less regular configuration or the maintenance of process extensions^61,62^ (Figure 5C). The upregulation of cell-cell adhesion molecules known to bind to each other (ITGAL, ICAM1) could be responsible for establishing contacts between microglia or attenuating repulsion^62^ (Figure 5C).

Finally, we looked at signaling pathways known to be regulated by CD72 in B cells. CD72 is an inhibitory coreceptor to the BCR^44,63^. Upon antigen binding to BCR, associated tyrosine kinases such as LYN phosphorylate the immunoreceptor tyrosine-based inhibitory motif (ITIM) on CD72, which then recruits the tyrosine phosphatase SHP-1, leading to downmodulation of BCR signaling, thereby regulating the balance of activation and inhibition to prevent over-stimulation of the B cell^64^. Although LYN and SHP-1 can work downstream of many cell-surface receptors, we observed all three (CD72, LYN, and SHP-1) proteins to be upregulated in microglia treated with rCD72, thus presenting a possible CD72 gain-of-function mechanism (Figure 5D). Furthermore, we observed changes in the mitogen-activated protein kinase (MAPK) pathway, a major BCR-mediated signaling pathway: ERK1/2 (both downregulated), a module of MAPK that mediates cell proliferation, growth, and differentiation, is known to be inhibited by CD72 signaling^63,65^ (Figure 5D).

These data taken together suggest that adding rCD72 to microglia acts as a gain of function of endogenous CD72 signaling, which might have an inhibitory coreceptor function to an unknown microglial receptor that is analogous to the BCR in B cells (Figure 5D). Other molecular changes induced by rCD72 treatment, namely altered adhesion, cytoskeletal reorganization, and altered repulsive receptor composition are consistent with our observed morphological disruption of tiling (Figure 5C).

### CD72 overexpression in microglia recapitulates aspects of rCD72-mediated tiling disruption

To test our hypothesis that rCD72 induces a gain of function of endogenous CD72 signaling, we utilized a custom AAV9-derived capsid to efficiently transduce microglia *in vitro* with either mouse CD72 cDNA sequence or a 3xFlag tag sequence under the control of the CAG promoter (Figure 6A). We confirmed overexpression of CD72 protein in CD72-overexpressing microglia relative to 3xFlag-overexpressing microglia (Figures 6B-C). Tiling analysis revealed CD72-overexpressing microglia to have significantly decreased NND, significantly increased area coverage, and a nonsignificant decrease in spatial regularity (p=0.17) (Figures 6D-E). These changes were consistent with effects observed from rCD72 treatment, supporting our gain-of-function hypothesis (Figure 3F). This suggests that endogenous CD72 primarily controls the spatial arrangement of microglia, where baseline levels of CD72 contribute to consistent, regular spacing, whereas increased levels of CD72 promote rearrangement to tighter spacing and/or irregularity. On the other hand, we observed no change in contacts or skeleton complexity relative to the 3xFlag control (Figures 6D-E), suggesting that the contacts and skeleton complexity effects observed with rCD72 treatment were a secondary or tertiary effect of the treatment. These results support our gain-of-function model and suggest that the proper regulation of CD72 levels is important for homeostatic tiling.

**Figure 6.**
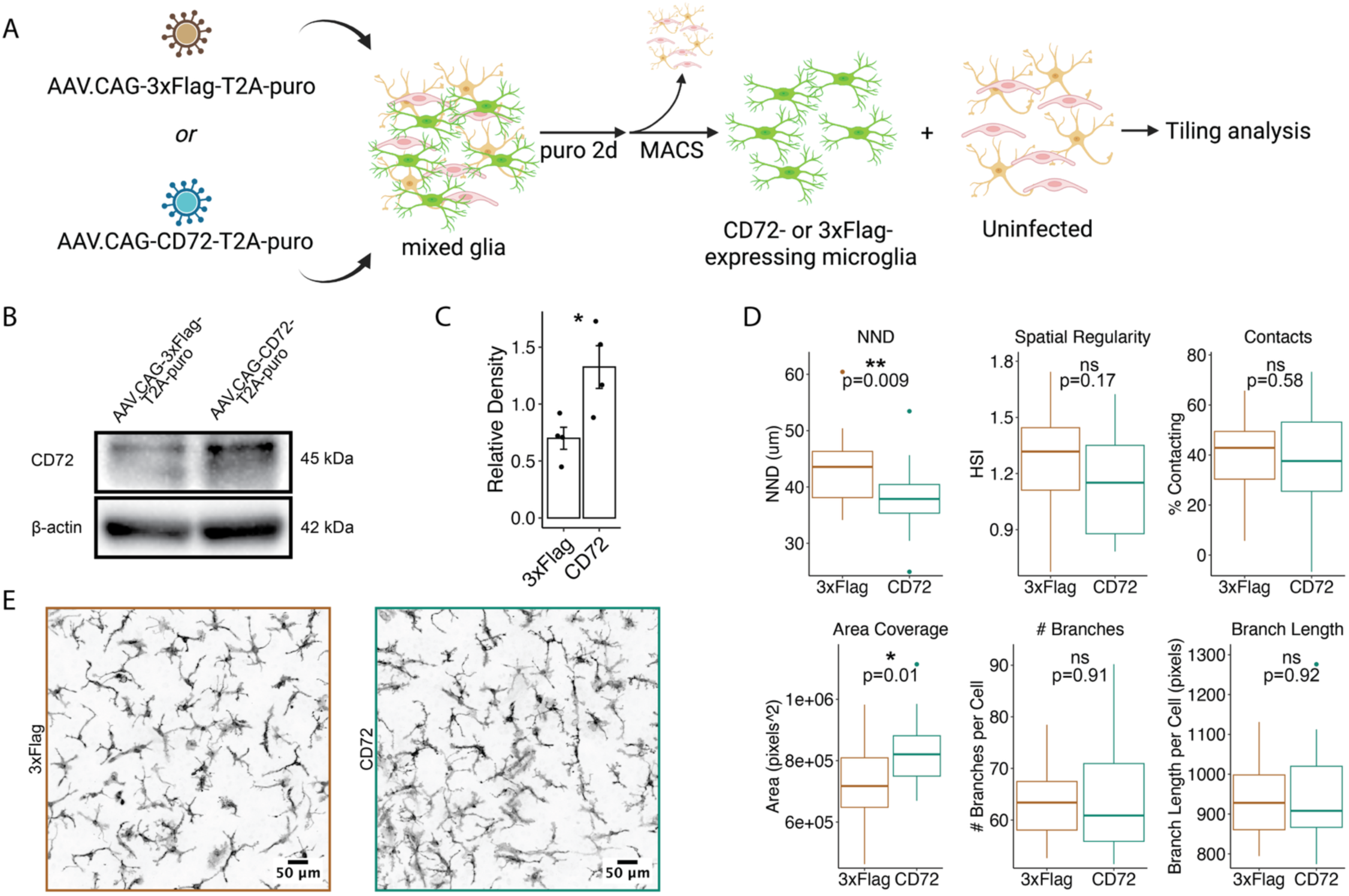
CD72 overexpression in microglia recapitulates aspects of rCD72-mediated tiling disruption. **(A)** Workflow describing AAV-mediated overexpression of CD72 (or 3xFlag as control) in mixed glia, puromycin (puro) selection, microglia enrichment via magnetic activated cell sorting (MACS), then replating with uninfected non-microglia. Tiling analysis was performed 48 hours later. (**B)** Western blotting analysis on lysate from puro-selected and enriched CD72- or 3xFlag-overexpressing microglia stained for CD72 and β-actin as loading control. **(C)** Quantification of four technical replicates from western blot in (B). CD72 bands are normalized to β-actin band densities. Bars show mean ± SEM. *p<0.05 by Welch’s T-test. **(D)** Quantifications of tiling features. N=four images in each of five wells, 20 technical replicates. All p-values by Welch’s T-test are listed. **(E)** Representative images of microglia over-expressing 3xFlag or CD72.

## DISCUSSION

Here, we present a formal study of microglial tiling and molecular mechanisms that regulate it. Assuming that tiling would be mediated by interactions at the cell surface, we employed a proximity labeling approach to the characterization of the microglial cell-surface proteome, which has previously only been performed in neurons^21,66,67^. Recognizing that the identified proteins could originate either from microglia or from other cell types that contact the microglial cell surface, we used scRNAseq to not only characterize the cell type-specific origins of these proteins, but also characterize a culture system that could be used for future studies of homeostatic microglial functions. We developed a screening method for functionally testing recombinant proteins’ effects on microglial tiling, characterized our lead candidate CD72 both *in vitro* and *in situ*, then used phosphoproteomics and genetic manipulation to propose a new potential molecular mechanism describing CD72’s role in the regulation of microglial spatial arrangement.

Although we focused on microglial tiling in homeostatic conditions, it stands to reason that similar mechanisms could apply in other conditions where spatial organization seems to matter. During initial colonization of the brain, microglia might require homotypic repulsion to establish tiling for the first time, as opposed to randomly settling in the brain parenchyma^10^. After microglia depletion via CSF1R inhibition, microglia clonally expand to repopulate the brain and proceed in a “wavefront” of proliferative cells towards microglia-free space, which could require active tiling mechanisms^68,69^. In addition, tiling is apparently disrupted in injury and disease, but the mechanism of tiling disruption is unknown, and it is not known how tiling regulation contributes to disease and/or recovery^15,16^. Thus, mechanisms of microglial sensing of neighbors (or lack thereof) and initiation of a response to regulate spatial distribution are a critical phenomenon for many neurobiological processes.

We established a foundation for the study of microglial tiling by formalizing a system for characterizing and quantifying microglial tiling; just as tiling is a complex and composite phenomenon, its measurement is a composite of the quantification of six features. We used this approach to screen the CSPs identified by mass spectrometry and found that CD72 is at least partially responsible for disrupting tiling by rearranging microglial spatial distribution. The function of CD72 in microglial is poorly described. In B cells, CD72 inhibits B cell activation constitutively and also upon antigen binding to the BCR, by its ITIM motif^64^. Microglia are not known to express the BCR, thus we hypothesize that CD72 acts as a regulatory coreceptor to a different microglial receptor; future work will aim to identify and characterize this receptor. Notably, SEMA4D is the only known ligand of CD72^70^. SEMA4D expressed by T cells can bind CD72 on B cells, which reduces phosphorylation of CD72’s ITIM motif, leading to reduced signal inhibition and enhanced B cell activation^64^. Based on this literature and our results showing that CD72 disrupts tiling, we would expect SEMA4D to promote tiling; however, we did not observe that effect in our screen, suggesting that the mechanisms of CD72 function in microglia vary from those that operate in B cells. Further investigation is necessary to confirm that there is a primary receptor in microglia that CD72 is acting alongside, identify this receptor, and identify any potential ligands.

CD72 was recently revealed as a pro-inflammatory factor in microglia in a mouse model of stroke: it is upregulated after stroke, and selective knockdown of CD72 in microglia reduced infarct volume, reduced inflammatory cytokine production, and reduced neuronal apoptosis^71^. This is consistent with our proteomic data showing pro-inflammatory immune response after rCD72 treatment and further exemplifies how tiling disruption and microglial activation seem to be inherently linked. Future studies should aim to determine whether 1) increased CD72 induces microglial activation and whether this leads to a disruption of tiling, 2) the disruption of tiling via increased CD72 is sufficient to induce microglial activation, or 3) if both are caused by CD72 via independent parallel mechanisms.

In conclusion, our study opens a new avenue of exploration into molecular mechanisms that regulate tiling, enabling the interrogation of questions about the biological meaning of tiling. This work is hoped to provide a foundation not only for the study of microglial tiling, but also for the study of cellular spatial organization across biology.

## RESOURCE AVAILABILITY

### Lead contact

Further information and requests for resources and reagents should be directed to the lead contact, Marius Wernig.

### Materials availability

No new unique materials were generated in this study.

### Data and code availability

All data reported in this paper will be shared by the lead contact upon request.

## ACKNOWLEDGEMENTS

We thank members of the Wernig lab, especially D. Wu, M. Mader, and Y. Shibuya for expertise and feedback on the project; PhD thesis committee members K. Shen, M. Buckwalter, B. Zuchero, T. Wyss-Coray for feedback on the project; T. Tsubata and T. Shichita for expertise on CD72; A. Yaron for advice on reagents; E. de la Serna for advice on image analysis; T. Sudhof, Y. Zhang, P. Lin, C. Vitale, Y. Ng for access to vibratome equipment; A. Lang and M. Vangipuram for administrative support; C. Caridi and the Stanford High-Throughput Screening Knowledge Center (supported by NIH 1S10OD026899-01); G. Wang and the Stanford Neuroscience Microscopy Service; the FACS core at the Stanford Institute for Stem Cell Biology and Regenerative Medicine. The work was supported by a Howard Hughes Medical Institute Faculty Scholar Award (M.W.), the Kleberg Foundation, and the California Institute for Regenerative Medicine (DISC0-13875). T.C.C was supported by a Neuroscience Research Training Grant (NIH T32MH020016) through the Stanford Neurosciences PhD program and a Stanford Interdisciplinary Graduate Fellowship affiliated with the Wu Tsai Neurosciences Institute. S.M.S. was supported by a Stanford Bio-X Fellowship. AAV generation and production was supported by R21MH126409 (B.E.D. and B.S.) and the Stanley Family Foundation. Cartoon elements of figures were created with BioRender.com.

## AUTHOR CONTRIBUTIONS

T.C.C. and M.W. were responsible for the conception, design, and interpretation of results and wrote the manuscript. T.C.C. performed all experiments and analysis except for what follows. M.R. performed optimization of genetic manipulation. M.X.W. assisted with genotyping, screen optimization, mass-spectrometry sample preparation, and image analysis code. S.Z. performed scRNAseq analysis. S.M.S. and T.W.C. performed label-free LC-MS/MS for cell-surface proteomics. S.R.S., J.P., and S.P.G performed TMT LC-MS/MS for microglial proteome and phosphoproteome. T.B., K.W., and D.K.V. cultured and treated organotypic hippocampal brain slices. M.A., O.H., and T.W.C. performed scRNAseq library preparation. S.J., B.S., and B.E.D. provided custom AAV capsid and produced virus. S.A.S. and L.L. provided iPEEL mice. A.N. assisted with mass-spectrometry sample preparation. Y.Y. assisted with FACS collection for scRNAseq and aligning reads. V.G.H. and M.S.B. provided bulk RNA sequencing data of microglia after stroke.

## STAR★METHODS

### Key Resources Table

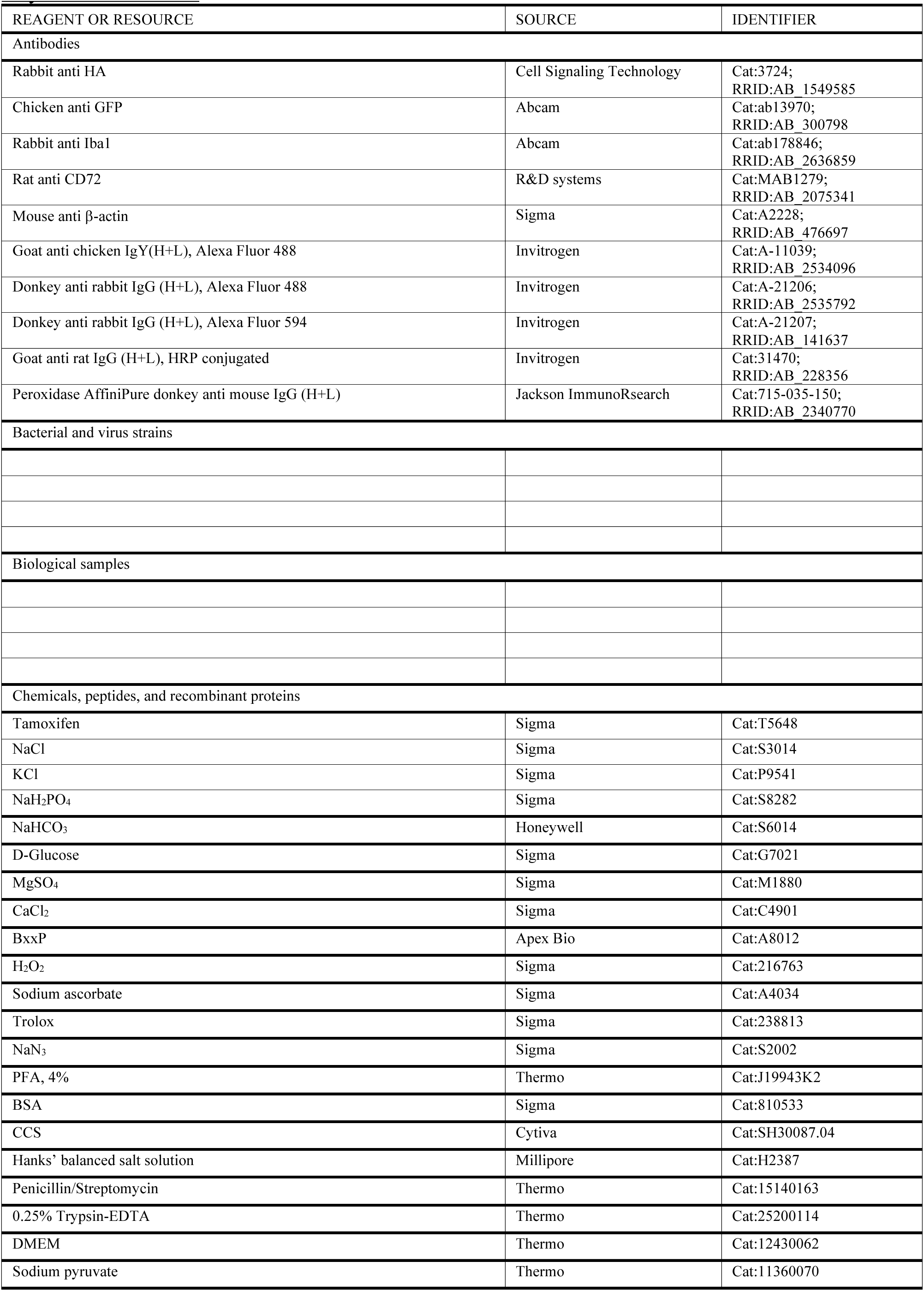
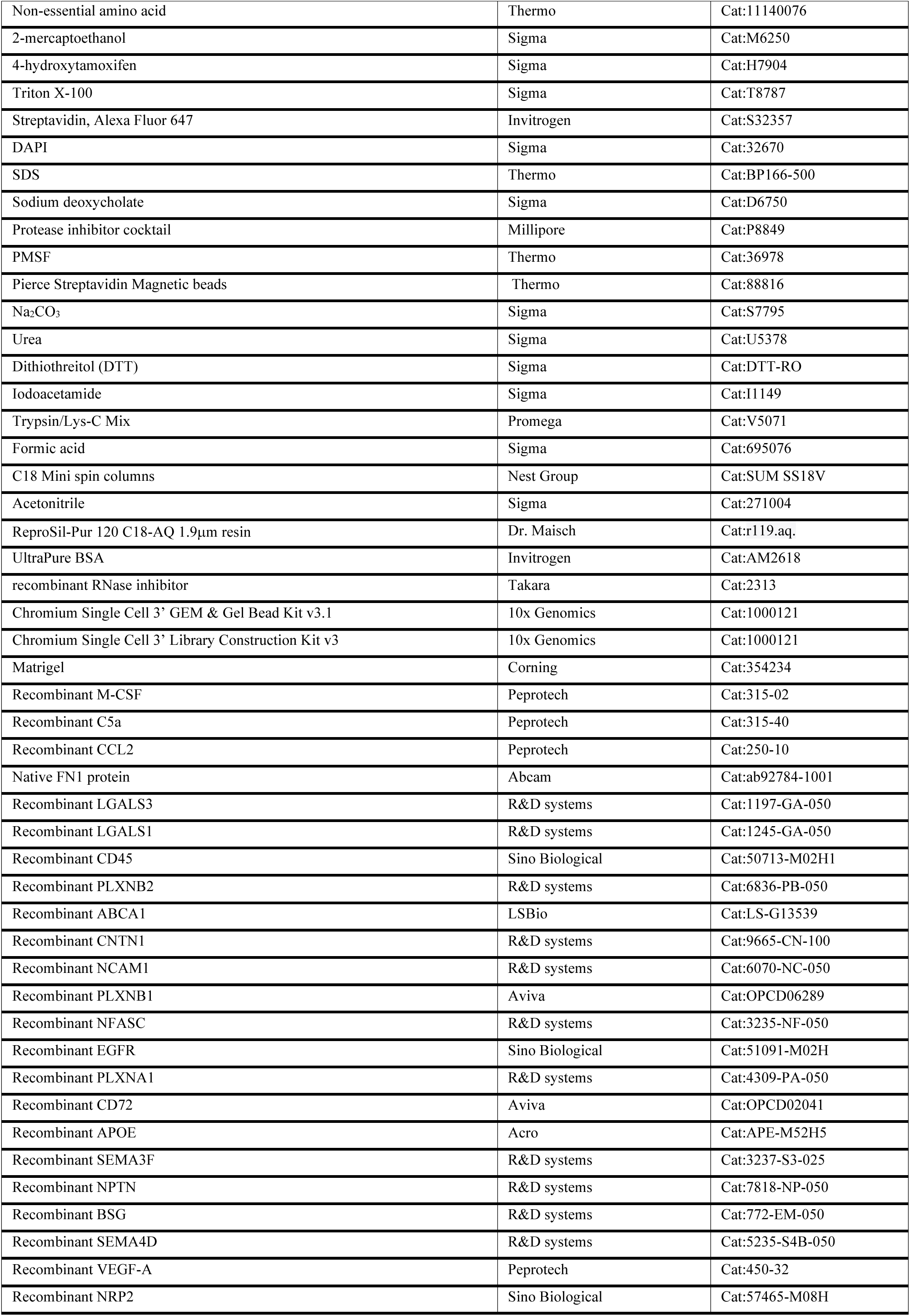
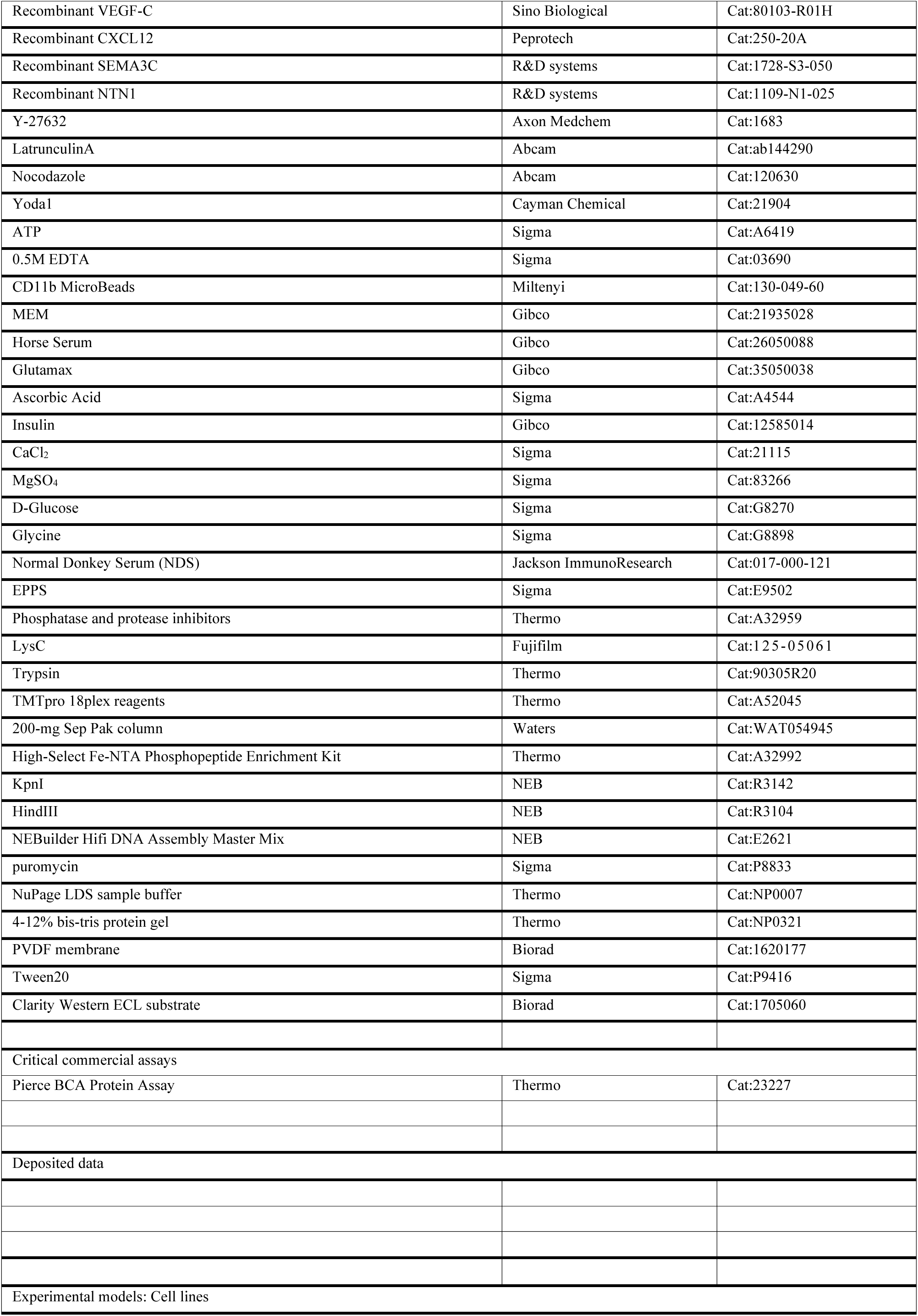
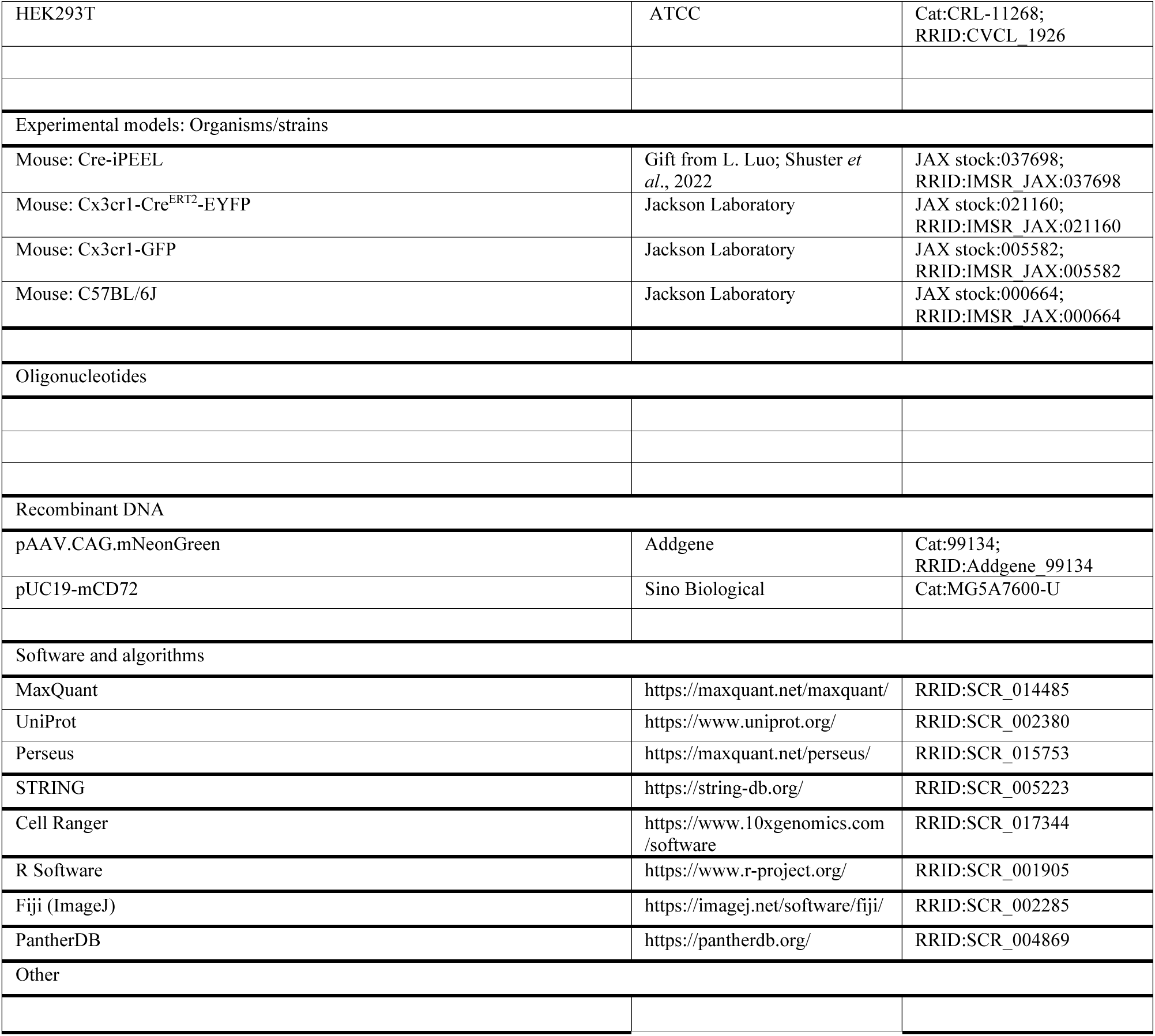

### Method Details Mice

All animal studies and procedures were approved by the administrative panel on laboratory animal care at Stanford University. All mice were group housed (up to five mice) under standard condition 12 h light/dark cycle with water and standard chow *ad libitum*. Mice used for proximity labeling were Cre-iPEEL, donated by Liqun Luo, crossed with Cx3cr1-Cre^ERT2^-EYFP, to achieve experimental mice homozygous for the Cre-dependent HRP allele and heterozygous for the Cx3cr1-Cre^ERT2^-EYFP. Littermate controls were also homozygous for the Cre-dependent HRP, but did not contain the Cx3cr1-Cre^ERT2^-EYFP. For screening, Cx3cr1-GFP mice were bred to C57BL/6J. Resulting litters, heterozygous for Cx3cr1^GFP^ allele, were sacrificed for primary mixed glia culture.

### Proximity labeling in acute brain slices

Cx3cr1-Cre^ERT2^-EYFP; iPEEL mice and littermate controls were injected intraperitoneally with 100mg/kg Tamoxifen twice with 48 hours in between at 6-8 weeks old. Two weeks later, mice were sacrificed quickly and the brain surgically removed and placed in ice cold carbogenated (5% CO_2_, 95% O_2_) artificial cerebrospinal fluid (ACSF) containing (mM): NaCl (119), KCl (2.5), NaH_2_PO_4_ (1), NaHCO_3_ (26), 11 D-Glucose (11), MgSO_4_ (1.3), CaCl_2_ (2.5). pH was adjusted to 7.2-7.4. 250µm sagittal slices were cut on a vibratome and allowed to recover in carbogenated ACSF at 32°C for 30 minutes. The ACSF was replaced by ACSF containing 100µM BxxP at 32°C for 60 minutes. Proximity labeling was initiated by adding 0.3% H_2_O_2_ to the BxxP-ACSF at 1:100, yielding a final 0.003% H_2_O_2_, and the container was swirled to diffuse the H_2_O_2_. After 3 minutes of incubation, the reaction was terminated by replacing the BxxP-ACSF with carbogenated quencher solution: ACSF containing 10mM sodium ascorbate, 5mM Trolox, and 10mM NaN_3_. The reaction was quenched 3 times for 5 minutes each time. Slices were fixed in 4% PFA in PBS at 4°C overnight with shaking for histological analysis.

### Mouse mixed glia isolation and culturing

Postnatal day 2 pups were anesthetized on ice and brains were surgically removed into cold Hanks’ balanced salt solution with penicillin-streptomycin (HBSS -ions +P/S). Meninges, olfactory bulb, and cerebellum were removed. Brain was transferred to clean HBSS -ions +P/S and minced using a clean #10 scalpel. Content was transferred to collection tube with 3ml cold HBSS -ions +P/S on ice. 1ml of 0.25% Trypsin-EDTA was added to each collection tube and incubated in a 37°C water bath for 30 minutes, with inversion every 10 minutes to mix. 2ml complete MEF media (DMEM with sodium pyruvate, non-essential amino acids, P/S, 10% CCS, 2-mercaptoethanol) was added to each tube to quench trypsin. The mixture was triturated with a 10ml serological pipette 20 times, or until chunks were broken down. Mixture was spun at 1000rpm for 3 minutes and supernatant was removed. The pellet was resuspended in complete MEF media and plated into a 10cm dish and incubated in a humidified 37°C, 5% CO_2_ incubator for about 5 days, changing the media every 3 days, until confluent. Cells were lifted using 0.25% Trypsin-EDTA and every 10cm plate was frozen into two cryogenic tubes (2 million cells). For all subsequent experiments, 2 million frozen glia were thawed quickly and cultured in 15cm plates using MEF media, changing media every 3 days, until confluent and microglia have grown in, 10-14 days. At this point, the mixed glia were ready for experiments in 15cm dishes, or for replating into appropriate vessels for other experimental set ups.

### Proximity labeling in cultured mixed mouse glia

2 million mixed glial cells isolated from Cx3cr1-Cre^ERT2^-EYFP; iPEEL and littermate control (iPEEL) mice were cultured and proximity labeled in 15cm dishes. N=3 of each of the following conditions were used: the experimental condition was Cx3cr1-Cre^ERT2^-EYFP; iPEEL glia treated with all labeling reagents, control #1 was Cx3cr1-Cre^ERT2^-EYFP; iPEEL glia treated with all labeling reagents except H_2_O_2_, control #2 was littermate control (iPEEL) glia treated with all labeling reagents. 4-hydroxytamoxifen was added to the MEF medium of all dishes at 1µM final concentration every other day for 4 days total. On the day of proximity labeling, media was removed and cells were washed once with warmed PBS. PBS was removed and replaced with warmed PBS containing 100µM BxxP and incubated at 37°C for 5 minutes. To the appropriate conditions to be treated with H_2_O_2_, 0.3% H_2_O_2_ was added at 1:100, yielding a final 0.003% H_2_O_2_. The dish was swirled for 45 seconds, then the solution was exchanged for quencher solution: PBS containing 10mM sodium ascorbate, 5mM Trolox, and 10mM NaN_3_. The reaction was quenched 3 times total for 3 minutes each at room temperature.

### Histological analysis on proximity labeled samples

For proximity-labeled acute brain slices, free-floating PFA-fixed slices were washed three times shaking in PBS at room temperature for 10 minutes each, blocked in blocking buffer (4% BSA, 1% CCS, 0.2% Triton X-100 in PBS) for 2 hours shaking at room temperature, incubated with primary antibodies rabbit anti HA 1:1000, chicken anti GFP 1:1000 in blocking buffer over two nights shaking at 4°C. The tissue was washed three times shaking in PBS at room temperature for 10 minutes each, then incubated with secondary antibodies goat anti chicken IgY Alexa Fluor 488 1:1000 and donkey anti rabbit IgG Alexa Fluor 594 1:1000, with the addition of Streptavidin 647 1:1000, in blocking buffer overnight shaking at 4°C. The tissue was washed once with PBS, once with DAPI in PBS, then twice with PBS, all shaking at room temperature for 10 minutes, mounted on glass slides and cover slipped, then imaged using a Zeiss LSM980 confocal microscope.

For proximity-labeled cultured mixed glia, quenched samples were fixed with 4% PFA at room temperature for 10 minutes. All washes starting here were half volume washes to reduce cell detachment. Cells were washed four times with PBS at room temperature for 5 minutes each. Cells were blocked in blocking buffer for 1 hour at room temperature, incubated with primary antibodies rabbit anti HA 1:1000, chicken anti GFP 1:1000 in blocking buffer overnight at 4°C, washed with PBS four times, incubated with secondary antibodies goat anti chicken IgY 488 1:1000 and donkey anti rabbit IgG 594 1:1000, with the addition of Streptavidin 647 1:1000 for 2 hours at room temperature, washed once with PBS, once with DAPI in PBS, then twice with PBS. Imaging was performed on a Leica DMi8 widefield microscope or Zeiss LSM980 confocal microscope.

For immunohistochemical analysis of cell-surface localization of biotin signal, samples were treated exactly as previous paragraph, while simply omitting Triton X-100 from all staining buffers.

### Cell lysis, streptavidin enrichment and mass spectrometry sample preparation (*in vitro* cell-surface proteomics)

Proximity labeled 15cm dishes were placed on ice and scraped thoroughly to lift the cells. Contents were transferred to a conical tube and triturated with a 10ml serological pipette 5 times. All tubes were centrifuged at 1000rpm for 3 minutes, supernatant was removed, and cell pellets were flash frozen in liquid nitrogen and stored at -80°C until further use.

Cell pellets were thawed and resuspended in 320µl of 1% SDS RIPA-Q buffer: 50 mM Tris-HCl [pH] 8.0, 150 mM NaCl, 1% SDS, 0.5% sodium deoxycholate, 1% Triton X-100, 1x protease inhibitor cocktail, 1mM PMSF, 10mM sodium azide, 10mM sodium ascorbate, 5mM Trolox. The samples were vortexed briefly, then probe sonicated at 4°C with 50% amplitude 2 times for 10 seconds each, with 1 second pause in between. The 1% SDS RIPA-Q buffer was diluted down to 0.2% SDS and samples rotated at 4°C for 15 minutes. Samples were centrifuged at 16,000g for 10 minutes at 4°C and supernatant was transferred to a new tube on ice. BCA assay was performed on the lysate to determine total protein concentration.

Streptavidin magnetic beads were washed twice with 0.2% RIPA-Q by rotating at room temperature. 320µl of beads were added to roughly 18.6mg of total protein per sample and the mixtures were gently rotated at 4°C overnight. The following day, beads were washed with 1 ml of each of the following for 3 minutes rotating at room temperature: twice with 0.2% SDS RIPA-Q, once with 1M KCl, once with 0.1M Na_2_CO_3_, once with 2M urea in 10mM Tris-HCl pH 8, twice with 0.2% SDS RIPA-Q buffer.

Beads were washed twice with 200µl of 50mM Tris HCl pH 8, twice with 200µl of 2M urea in 50mM Tris HCl pH 8, then the wash buffer was completely replaced with 80µl of 2M urea in 50mM Tris HCl pH 8 containing 1mM dithiothreitol (DTT) and 0.4µg Trypsin/Lys-C. The samples were incubated for 1 hour at 25°C with 1000rpm shaking. The tubes were flicked every 15 minutes to fully resuspend the beads. The supernatant was moved to a new tube. Remaining beads were washed twice with 60µl of 2M urea in 50mM Tris HCl pH 8, each wash was combined with the supernatant. The resulting supernatant with digested peptides underwent reduction with 4mM DTT for 30 minutes at 25°C shaking 1000rpm, then alkylation with 10mM iodoacetamide for 45 minutes in the dark at 25°C shaking 1000rpm. An additional 0.5µg of Trypsin/Lys-C was added for overnight digestion at 25°C shaking 700rpm. The samples were acidified to pH < 3 by adding formic acid to 1% of the final volume. Samples were desalted using C18 Mini spin columns, eluted with 50µl of 80% acetonitrile/1% formic acid, then dried to completion.

### Label-free liquid chromatography-tandem mass spectrometry (*in vitro* cell-surface proteomics)

Samples were analyzed by online nanoflow LC-MS/MS using a Q Exactive HF-X (Thermo Fisher Scientific) coupled with an UltiMate 3000 RSLCnano system (Thermo Fisher Scientific). Peptides were reconstituted in 10 μL of 0.1% formic acid in HPLC-grade water. 2 μL of each sample were loaded on an in-house 75 μm (inner diameter) capillary column packed with 40 cm of ReproSil-Pur 120 C18-AQ 1.9μm resin. Chromatographic separation was achieved using a flow rate of 0.3 μL/min with the following 120 min gradient: 96% A + 4% B for 18 min, 70% A + 30% B for 72 min, 60% A + 40% B for 15 min, and 4% A + 96% B for 15 min, where solvent A was 0.1 % formic acid in HPLC-grade water and solvent B was 0.1% formic acid in HPLC-grade acetonitrile. Full MS scans were acquired at a resolution of 60,000, with an automatic gain control (AGC) target of 3e6, maximum injection time (IT) of 20 ms, and scan range 300–1650 m/z in a data-dependent mode. MS2 scans were acquired with the following parameters: resolution of 15,000, AGC target of 1e5, maximum IT of 54 ms, loop count 15, TopN 15, isolation window 1.4 m/z, fixed first mass 100.0 m/z, normalized collision energy (NCE) 28 units, charge exclusion of unassigned, 1, 6-8, and >8, peptide match preferred, exclude isotopes on, and fragmented m/z values were dynamically excluded from further selection for a period of 45 s. Raw data were processed using MaxQuant software (version 1.6.10.43)^72^. Peptide spectral matches were made against a target-decoy Mus musculus reference proteome database from Uniprot. Methionine oxidation and N-terminal acetylation were specified as variable modifications, and carbamidomethylation of cysteines was specified as a fixed modification. Precursor ion search tolerance of 20 ppm and product ion mass tolerance of 20 ppm were used for searches. Both unique and razor peptides were used for quantification. Results were filtered to a 1% FDR at the peptide and protein levels. Proteins were quantified and normalized using MaxLFQ^73^ with a label-free quantification (LFQ) minimum ratio count set to 1.

### Proteomics data analysis (*in vitro* cell-surface proteomics)

Using Perseus software, 2064 detected protein groups were filtered by reverse, contaminants, only identified by site, minimum 2 valid values in each condition, and missing values were imputed^74^. Ratiometric analysis was applied as performed previously^21,66,75^. In short, proteins were categorized according to their gene ontology (GO) Cellular Component and Subcellular Localization annotations downloaded from the UniProt database (retrieved March 2022). Proteins that were annotated as “plasma membrane” in the GO Cellular Compartment or “secreted” in Subcellular Location were categorized as true positives (TPs), for which there were 712.

Proteins that did not have either of those annotations, but did have “nucleus”, “mitochondria”, or “cytoplasm” in the GO Cellular Compartment were categorized as false positives (FPs), for which there were 1,041. For each protein, the average LFQ intensity from three replicates in each condition was calculated, then two ratios of LFQ intensities were calculated: Exp / Control 1 and Exp / Control 2. The proteins were ranked in descending order according to these ratios. The top 150 highest ratios from each list were uploaded to the STRING database (v11.5) search portal and significant enriched GO Cellular Compartments were plotted to check the enrichment for CSPs. The search was against the mouse reference proteome as the background. For each protein on each ranked list, the accumulated true- and false-positive counts above its ratio were calculated. A receiver operating characteristic (ROC) curve was plotted for each ratio. The cutoff was set where the true-positive rate minus false-positive rate (TPR – FPR) was maximized: for Exp / Control 1 the cutoff was 0.222, for Exp / Control 2 the cutoff was 0.635. Post-cutoff proteomic lists were intersected to obtain the final proteome. For GO analysis of the final proteome of 241 proteins, we uploaded the proteome to the STRING database search portal and plotted the top enriched GO Biological Processes with the lowest false discovery rates. The search was against the mouse reference proteome as the background.

### scRNAseq sample preparation

scRNAseq on mixed mouse glia isolated from littermate control iPEEL pups (homozygous for the Cre-gated HRP gene, but lacking Cx3cr1-Cre^ERT2^-EYFP) was performed. Cells were washed once with PBS, dissociated with 0.25% Trypsin-EDTA, quenched with MEF media, then collected and centrifuged at 1000rpm for 3 minutes. All subsequent steps were performed on ice. The cell pellet was resuspended in FACS buffer—PBS with 1% UltraPure BSA and recombinant RNase inhibitor. DAPI was added shortly before 100,000 live cells were sorted on a BD FACSAria II Cell Sorter. Collected cells were centrifuged at 400 x g for 5 minutes at 4°C with break 2. Supernatant was removed leaving 40 ul suspended cells. Cells were counted and assessed for concentration and quality.

### scRNAseq library construction

Reagents of the Chromium Single Cell 3’ GEM & Gel Bead Kit v3.1 were thawed and prepared according to the manufacturer’s protocol. Cells and master mix solution was adjusted to target 10,000 cells per sample and loaded on a standard Chromium Controller (10X Genomics) according to manufacturer protocols. Library construction was conducted using Chromium Single Cell 3’ Library Construction Kit v3. All reaction and quality control steps were carried out according to the manufacturer’s protocol and with recommended reagents, consumables, and instruments. 11 PCR cycles were applied for library generation. Quality control of cDNA and libraries was conducted using a Bioanalyzer (Agilent) at the Stanford Protein and Nucleic Acid Facility.

Illumina sequencing of the resulting libraries was performed by Novogene (https://en.novogene.com/) on an Illumina NovaSeq S4 (Illumina), sequenced to 20K paired reads per cell. Base calling, demultiplexing, and generation of FastQ files were conducted by Novogene. The 10× Genomics Cell Ranger (v.7.1.0) analysis pipelines were used to align reads to the mm10 reference genome.

### scRNAseq data quality control and analysis

The sequencing data was imported into R (v4.4.0) to be analyzed. The original dataset was composed of 13056 cells with 21055 genes or features. Doublets were removed using scDblFinder tool (v1.14.0)^76^. scDblFinder was applied across samples with a deviation from the input doublet rate (dbr.sd) set to 0.1, balancing the identification of true doublets with the retention of genuine singlet cells, leaving 11672 cells for a total of 21055 genes. Seurat (v5) was employed to filter for cells with RNA molecule counts (nCount_RNA) between 2,000 and 20,000, cells expressing more than 800 unique genes (nFeature_RNA), cells with mitochondrial RNA content (percent.mt) less than 15%, and cells with hemoglobin gene expression (percent.hb) less than 0.1%^77^.

These numbers were determined by looking at the overall map of the normal distribution of the cells across these variables and determining what cutoffs would define cells of lower quality that could skew downstream analysis. Furthermore, we eliminated genes expressed in fewer than 10 of the remaining cells. After all filtering, the data comprised 11126 cells and 18075 genes. SCTransform normalization was applied and principal component analysis (RunPCA()) on the top 2,000 variable genes facilitated dimensionality reduction, with the top 20 principal components selected using the elbow plot heuristic. Construction of a k-nearest-neighbors graph through the FindNeighbors() function enabled robust neighborhood-based analysis. Unsupervised clustering, employing the Louvain algorithm via the FindClusters() function at a resolution of .15, delineated 5 distinct clusters. This lower resolution was used to primarily delineate microglia and non-microglia.

Dimensionality reduction to two-dimensional space, achieved via uniform manifold approximation and projection (UMAP) on the top 20 principal components using the RunUMAP() function, facilitated intuitive visualization of cellular relationships. FindAllMarkers() function facilitated identification of DEG markers for each cluster and labelling of the 5 clusters as non-microglia or microglia. Then, a second round of normalization and clustering of subsetted microglia and non-microglia was performed using the same processing steps. The DEGs were again extracted and used to identify cell types using canonical markers found through various literature. DoHeatmap(), FeaturePlot(), and DotPlot(), were used to generate figures for visualization.

### Recombinant protein screen in mixed glia culture

All purchased recombinant proteins were dissolved in 0.1% BSA in PBS to a stock concentration of 50µg/ml and stored at -80°C until use. To prepare the screening plate, Matrigel was diluted 1:50 in DMEM on ice and working concentration of all recombinant proteins were made up in the diluted Matrigel mixture to a final 400ng/ml. The vehicle control was 0.1% BSA, with the same volume added to the Matrigel mixture as all other recombinant protein aliquots to make the working concentration, resulting in a final 0.0008% BSA, or 8ug/ml BSA. Working concentrations for other agents were: Y-27632 10µM, LatrunculinA 100nM, Nocodazole 100nM, Yoda1 500nM, ATP 600µM. Working dilutions of all treatments in Matrigel mixture were distributed in triplicate to a 96-well culture plate on ice, then incubated in a humidified 37°C, 5% CO_2_ incubator for 1 hour.

For the soluble version of the screen, all recombinant proteins and other agents were treated to the same concentrations, but in the cell culture media after cell plating, instead of embedded in Matrigel prior to cell plating.

Cx3cr1-GFP mixed glia were cultured according to above section (<u>Mouse mixed glia isolation and culturing</u>). On the day of cell replating, glia were washed once with with PBS, dissociated with 0.25% Trypsin-EDTA, quenched with MEF media, then collected and centrifuged at 1000rpm for 3 minutes. All subsequent steps were performed on ice with cold media. Cells were resuspended to singe cell suspension in PBS, counted, centrifuged again, PBS was removed. Cells were resuspended in 80ul MACS buffer – 0.5% BSA, 0.4% 0.5M EDTA in PBS – per 10^7^ total cells and stained with 20ul CD11b MicroBeads for 15 minutes at 4°C. Cells were washed by adding 2ml of MACS buffer, centrifuging 1000rpm for 3 minutes, and discarding the supernatant. Cells were resuspended in 500ul of MACS buffer and separated on a magnetic QuadroMACS Separator (Miltenyi) according to the manufacturer’s protocol. Enriched (microglia) and flow-through (non-microglia) cells were counted separately, then recombined at ∼15% microglia mixture in MEF media. The 96-well culture plate containing Matrigel-embedded screen candidates was removed from the incubator and all liquid was removed. The ∼15% microglia cell mixture was distributed to the 96-well culture plate at 50K total cells per well (7,500 microglia; 42,500 non-microglia) and returned to the incubator for 2 days.

For the soluble version of the screen, recombinant proteins were added into the media after cell plating.

### Fixation and immunofluorescent staining of recombinant protein screen

All wells were washed once with PBS, then fixed with 4% PFA for 10 minutes at room temperature. Half volume washes were performed using PBS 5 times for 5 minutes each. Cells were blocked with blocking buffer – 4% BSA, 1% CCS, 0.2% Triton X-100 in PBS for 1 hour at room temperature. Buffer was exchanged for blocking buffer containing chicken anti-GFP primary antibody at 1:1000 for overnight incubation at 4°C. Half volume washes were performed using PBS 5 times for 5 minutes each. Buffer was exchanged for blocking buffer containing goat anti-chicken IgY Alexa Fluor 488 secondary antibody at 1:1000 for 1 hour at room temperature. Half volume washes were performed using PBS 5 times for 5 minutes each. On the 3^rd^ wash, DAPI was added to the PBS to final 1:50,000.

### Automated imaging of recombinant protein screen

High-content (automated) confocal imaging was performed on an ImageXpress Micro Confocal Microscope (Molecular Devices). 50 µm slit confocal mode was used with 20X 0.75 Plan Apo 1mm WD objective in both DAPI and FITC filters. Each image projected through 13µm of z thickness at 1µm step size and the maximum intensity projection was saved as the raw image. 4 sites near the center of the well were imaged for each well (triplicates), resulting in N=12 for each candidate.

### Image processing and analysis of tiling features

Fiji (ImageJ version 2.14.0/1.54f) was used for all batch thresholding and quantifications. Macros were written for all parts of the image processing, and R was used for analysis and data visualization.

To calculate NNDs, the (x,y) coordinates of each microglial nucleus was first acquired. DAPI and GFP images were thresholded using Huang method and regions of interest (ROIs) were created for each image.

DAPI thresholded images were Watershed. The two corresponding ROIs for each image were overlapped using the “AND” function within ROI manager to only retain the nuclei overlapping with microglia. Fiji’s Analyze Particles function was run on the retained nuclei with size cutoff 450 – infinity. Separate result files for each image containing the (x,y) coordinates of the centroid of each nucleus within that image were saved. In R, the spatstat package was used to convert the points into a ppp object with the dimensions of the image and calculate the NND for each point^78^.

To measure spatial regularity, the Hopkins-Skellam test of complete spatial randomness was performed on the same ppp object using the spatstat package^78,79^. The test statistic (Hopkins-Skellam Index) was used as a simple summary of spatial clustering, randomness, or regularity^13^.

To acquire area coverage data, Analyze Particles was run solely on the GFP thresholded images using size cutoff 450 – infinity and separate results files for each image containing all particle areas were saved. In R, every result file was summed for the total pixel area of that image.

Contacts measures the degree of physical contact between microglia and is measured by: the number of microglial nuclei – the number of particles observed by Fiji after thresholding the image only in the GFP channel, divided by number of microglial nuclei. This relies on the fact that the Analyze Particles tool in Fiji segments continuous signal as a single observed object, thus microglia that are in physical contact must consequently be counted as a single observed object. If the true number of microglia is higher than the observed count, the difference represents some degree of contact. To get the true count, Analyze Particles was re-run on the GFP/DAPI overlapping nuclei while excluding signal on the edges of the image, with the same size cutoff (450 -infinity). To get the observed count, Analyze Particles was run on the GFP thresholded signal excluding signal on the edges. In R, the true count of microglia – the observed count was calculated for every image, then divided by the true count.

To acquire skeleton morphology data, the GFP thresholded images were skeletonized and analyzed by the “Analyze Skeleton” plug-in in Fiji. Results tables containing branch information were saved separately for each image. In R, low quality skeletons resulting from nonspecific puncta were filtered out by specifying # Junctions > 0 and # Branches >1. To calculate number of branches per cell, the number of branches in each image were summed, then divided by the true count of microglia, not excluding edges (total number of microglial nuclei counted from the NND analysis). To calculate branch length per cell, all branch lengths in each image were summed, then divided by the same true count of microglia.

P values were calculated using Welch’s t-test of every candidate against control within round. All P values from the experiment were collected and corrected for multiple hypothesis testing using Benjamini-Hochberg (BH) adjustment.

### Live confocal imaging of mixed glia and tiling analysis

For experiments where live (non-fixed) imaging was required (Figure 4C), Cx3cr1-GFP mixed glia were plated and treated solubly with candidate protein or control (vehicle 0.1% BSA). Image processing and analyses were performed differently from the fixed imaging screen—only using the endogenous Cx3cr1-GFP signal (no nuclear stain). We did attempt to use a Hoescht stain for live imaging, however, we found it to be toxic to cells. Imaging was performed on an ImageXpress Micro Confocal Microscope. 50 µm slit confocal mode was used with 20X 0.75 Plan Apo 1mm WD objective in the FITC filter. Imaging sessions on different days always returned to the same field of view within each well and all imaging parameters were the same across days. Each image projected through 13µm of z thickness at 1µm step size and the maximum intensity projection was saved as the raw image. 4 sites near the center of the well were imaged for each of 3 wells per condition, resulting in N=12. In Fiji, GFP images were thresholded using Triangle method, then Analyze Particles was applied using size cutoff of 350-infinity pixels. Result tables for each image were saved. In R, the average particle area within each image was calculated. The spatstat package was used on the (x,y) coordinates of every particle’s centroid to calculate the Hopkins-Skellam Index of spatial regularity for each image.

### Timelapse imaging and repulsion analysis

Timelapse imaging was performed on live Cx3cr1-GFP mixed glia on top of Matrigel embedded with candidate protein or control (vehicle 0.1% BSA) (see above: <u>Recombinant protein screen in mixed glia culture</u>). After two days of incubation, Cx3cr1-GFP signal was imaged live using the ImageXpress Micro Confocal Microscope. 50 µm slit confocal mode was used with 20X 0.75 Plan Apo 1mm WD objective in the FITC filter for 46 timepoints with ∼140s intervals between, totaling ∼105 minutes. One well of each treatment was imaged, two sites within each well were quantified, and 22-23 Rectangular ROIs were drawn based on the apparent contact event between two microglial cells at any timepoint. The ROIs were cropped and thresholded using Triangle method. Every time frame that had a continuous signal between the two cells (adjacent connecting or diagonally touching pixels) was recorded as contact. The number of time frames in which contact was occurring was divided by the total number of timepoints then divided by 100 to calculate the percent time contacting.

### Organotypic hippocampal slice culture preparation and recombinant protein treatment

C57BL/6J mice were used for the generation of organotypic hippocampal brain slice cultures (BSCs) in accordance with German Animal Protection Laws and registered as N 03/24 M. The preparation of BSCs was based on the protocol by Stoppini *et al*^80^. In short, pups were sacrificed on postnatal day 5 by decapitation with subsequent removal of the whole brain through careful cross-wise opening of skin and skull. The brain was transferred into cold preparation buffer (1x minimum essential medium (MEM), 2 mM Glutamax, pH 7.35) and both hippocampi dissected out. Hippocampi were cut into 350 µm thick slices using a tissue chopper (McIlwain), collected in fresh preparation buffer, and visually checked for preservation of slice integrity. 3 to 4 BSCs each were transferred onto sterile Millicell Cell Culture Inserts (Merck) in a 6-well plate with 1.2 ml of pre-warmed culture medium MEM + 20 % Horse Serum, 1 mM Glutamax, 0.00125 % Ascorbic Acid, 0.001 mg/ml Insulin, 1 mM CaCl2, 2 mM MgSO4, 13 mM D-Glucose, 100 U/ml/100 μg/ml Penicillin/Streptomycin; pH 7.28). Medium change was carried out three times a week and cultures kept at 37°C with 5% CO_2_. After 10 days, culture medium was removed, wells washed with PBS, and fresh medium supplemented with final concentrations 800 ng/ml rCD72 or control (vehicle 0.1% BSA) was added. Cultures were incubated for 4 days.

### Fixation and immunofluorescent staining of organotypic hippocampal slices treated with recombinant protein

Slices were fixed 4 days after treatment by briefly washing with PBS then submerged in 4% PFA overnight at 4°C while remaining on the inserts. Fixed slices were washed three times with PBS+100mM Glycine for 10 minutes each, blocked in 5% blocking buffer (5% NDS, 0.3% Triton X-100, 100mM Glycine in PBS), and incubated with primary antibody rabbit anti Iba1 1:1000 in 2% blocking buffer for 2 nights shaking at 4°C. The tissue was washed three times shaking in PBS+100mM Glycine at room temperature for 20 minutes each, then incubated with secondary antibody donkey anti rabbit IgG Alexa Fluor 488 1:1000 in 1% blocking buffer for 2 hours shaking at room temperature in the dark. The tissue was washed once with PBS+100mM Glycine, once with DAPI in PBS+100mM Glycine, then twice with PBS+100mM Glycine, all shaking at room temperature for 20 minutes in the dark. The tissue was mounted on glass slides and cover slipped, then imaged using a LSM980 confocal microscope (Zeiss) at 20x objective through 6 z-stacks of step size 2.75µm, then maximum intensity projected. For each treatment, 9 fields of view were imaged across 3 biological replicates.

### Tiling analysis in *in situ* organotypic hippocampal slice tissue

Maximum intensity projections were cleaned up via Subtract Background and Despeckle functions in Fiji. Iba1 images were thresholded using Huang method. Due to variability of signal depth within the tissue thickness, all tiling analyses were performed solely on maximum intensity projected Iba1 signal without contribution from the DAPI nuclear stain. Analyze Particles was applied to the Iba1 thresholded images with size cutoff of 50-infinity and results tables were saved for each image. (x,y) coordinates of each particle were used to determine NND and Hopkins-Skellam Index, the sum of all particles’ area within image was calculated for area coverage, and skeletons with # Junctions > 5 were used to quantify branch length (see details in section: <u>Image processing and</u> <u>analysis of tiling features</u>).

### Mass spectrometry-based proteomic and phosphoproteomic sample prep and TMT labeling (*in vitro* rCD72-treated microglia)

C57Bl6 mixed glia were cultured in a 15cm dish as described (<u>Mouse mixed glia isolation and culturing)</u>. 6 well culture plates were coated with Matrigel, embedded with rCD72 or control (vehicle 0.1% BSA) as described (<u>Recombinant protein screen in mixed glia culture</u>). Mixed glia were replated (without MACS enrichment) into the 6 well plates and incubated for 2 days with the treatments, then microglia were enriched out via MACS as described (<u>Recombinant protein screen in mixed glia culture</u>), cell pellets were flash frozen in LN, then stored at -80°C until further use.

A modified version of the SL-TMT protocol was followed^81^. To microglial cell pellets were added 200 µl lysis buffer (200 mM EPPS, pH = 8.5 with 8M urea, phosphatase and protease inhibitors, and 1 mM sodium pervanadate). Each pellet was lysed via probe sonication for 5s at 50% amplitude. After protein concentration assay via BCA, 100 µg of each lysate was dispensed and volume was increased to 100 µl (using lysis buffer) for each lysate with concentration < 1 µg/µl. Lysates were reduced with 5 mM DTT for 30 min at 37 °C, alkylated with 10 mM iodoacetamide for 30 min at room temperature in the dark, and then quenched with 10 mM DTT for 15 min at room temperature. Chloroform-methanol precipitation was then performed at a ratio of 4:1:4 methanol:chloroform:aqueous buffer (lysate + water) at a total volume of 900 µl by vortexing and then centrifuging at 17,000 rcf for 15 min at room temperature After removal of aqueous and organic phases, protein precipitates were washed in 400 µl methanol, the methanol was removed after centrifuging at 17,000 rcf for 30 min at room temperature, and then the proteins were resuspended in 100 µl 200 mM EPPS, pH = 8.5. Proteins were digested for 14 h with LysC (1:100 E:S ratio) and then 6 h with trypsin (1:100 E:S ratio). Peptides were then labeled with TMTpro 18plex reagents according to the manufacturer’s protocol. After assessment via “label check” analysis, the TMTpro reagents were quenched with 0.3% hydroxylamine for 15 min at room temperature and then the channels were combined accordingly. The resulting TMT plex was then desalted with a 200-mg Sep Pak column, evaporated in a vacuum centrifuge, and then subjected to phosphopeptide enrichment using the High-Select Fe-NTA Phosphopeptide Enrichment Kit (Thermo Fisher Scientific) according to the manufacturer’s protocol.

The phosphopeptides were desalted via StageTip^82^. Four layers Empore material (3M) were washed with 50 µl methanol then 50 µl 70% acetonitrile + 1% formic acid in water and then equilibrated with 2x 50 µl 3% acetonitrile + 1% formic acid in water by centrifuging at 1000 rcf for 3 min. Phosphopeptides were then resuspended in 100 µl 3% acetonitrile + 1% formic acid in water and loaded onto the StageTip. The buffer was then centrifuged through the StageTip and then the phosphopeptides were washed with 2x 50 µl 3% acetonitrile + 1% formic acid before elution with 2x 70% acetonitrile + 1% water into a mass spectrometry sample vial insert. The phosphopeptides were then evaporated in a vacuum centrifuge and then resuspended in 20 µl 3% acetonitrile + 1% formic acid in water.

Non-phosphopeptides were desalted with a 200-mg Sep Pak column using the same buffer sequence as above with StageTip desalting, except that a vacuum manifold was used to move buffer and 2.5 ml of buffer was used per wash and the peptides were eluted with 2x 600 µl 70% acetonitrile + 1% formic acid. After evaporation in a vacuum centrifuge, 300 µg of non-phosphopeptides were then fractionated via basic-pH high-pressure liquid chromatography (HPLC) by subjecting them to a 50-min linear gradient from 13% buffer B (90% acetonitrile in 10 mM ammonium bicarbonate) in buffer A (5% ACN in 10 mM ammonium bicarbonate) to 42% buffer B in buffer A at 0.6 ml/min on an Agilent 300Extend C18 column (3.5 µm particles, 4.6 mm ID, 250 mm length). 96 fractions were collected and then combined into 24 fractions as described in Navarrete-Perea et al., desalted via StageTip as described above, and then each fraction was resuspended in 12 µl 3% acetonitrile + 1% formic acid in water^81^.

### Multiplexed liquid chromatography-tandem mass spectrometry of proteome and phosphoproteome (*in vitro* rCD72-treated microglia)

Phosphopeptides were analyzed on an Orbitrap Astral mass spectrometer (Thermo Fisher Scientific) coupled with Neo Vanquish liquid chromatograph and equipped with a FAIMSpro interface (Thermo Fisher Scientific). Peptides were separated on a 110 cm µPAC C18 column (Thermo Fisher Scientific). For each analysis, we loaded ∼0.5 μg phosphopeptides onto the column. Peptides were separated using a 120 min gradient of 5 to 29% acetonitrile in 0.125% formic acid with a flow rate of 300 nL/min. The scan sequence began with an MS1 spectrum (Orbitrap analysis, resolution 180,000, 350-1650 Th, automatic gain control (AGC) target set to 200%, maximum injection time set to 50ms). The hrMS2 stage consisted of fragmentation by higher-energy collision-induced dissociation (HCD, normalized collision energy 35%) and analysis using the Astral (AGC 100%, maximum injection time 30ms, isolation window 0.7 Th, TMT mode was turned “on”). Data were acquired using the FAIMSpro interface in which the dispersion voltage (DV) was set to 5,000V. Dynamic exclusion was set to 15 s with a 5-ppm mass tolerance. The sample was analyzed twice with two sets of CVs: Set 1 = -35,-45,-55,-65 and -75V and Set 2 = -30, -40, -50, -60 and -70V. The dynamic exclusion list was shared across CVs. The TopN parameter was set at 25 scans per CV.

Non-phosphopeptide fractions were analyzed on an Orbitrap Eclipse mass spectrometer (Thermo Fisher Scientific) equipped with an EASY-nLC 1200 liquid chromatograph (Thermo Fisher Scientific) bearing a 30-cm column with 100-µm inner diameter packed with ReproSil-Pur C18-coated beads of 2.4-µm diameter (Dr. Maisch GmbH r124.aq.0001) according to the FlashPack protocol^83^. During a 90-min method, 1 µl of peptides were injected and then equilibrated for 5 min with 5–6% buffer B (95% acetonitrile + 0.1% formic acid in water) in buffer A (5% acetonitrile + 0.1% formic acid in water) and then separated using a 71-min linear gradient from 6% to 30% buffer B in buffer A, followed by a 7-min linear gradient from 30% to 50% buffer B, and then a 5-min gradient from 50% to 100% buffer B followed by 2 min of isocratic flow at 100% buffer B. A real-time search (RTS)-synchronous precursor selection (SPS)-MS3 method was used with the following parameters: for MS1, the orbitrap was operated at resolution = 120k with a scan range of 350–1500 Th and RF lens at 30% amplitude. The standard automatic gain control (AGC) feature was used, as was automatic maximum injection time (IT) calculation. All mass spectra were acquired in centroid mode with positive polarity. The monoisotopic peak determination feature was enabled; peptide-like ions with intensity > 5.0e3 and charge state in [2,5] were selected for MS2. The dynamic exclusion duration was 90 s with a mass tolerance of ±10 ppm; all isotopes and charge states were excluded. Precursors were isolated for MS2 analysis with an isolation window of 0.5 Th and then fragmented in the linear ion trap using collision-induced dissociation (CID) with a collision energy of 35% for 10 ms and a Q parameter of 0.25. MS2 fragments were analyzed in the linear ion trap at 66,666 Th/s with the automatic scan range and AGC target features enabled with a maximum IT of 35 ms. Real-time search was performed against the mouse reference proteome downloaded form Uniprot on October 3, 2024 using trypsin digestion including Lys-Pro cleavages. Cysteine carbamidomethylation (57.0215 Da) and lysine and peptide N-terminus TMTpro modification (304.2071 Da) were included as static modifications and methionine oxidation (15.9949 Da) was included as a variable modification. Two missed cleavages and two variable modifications per peptide were allowed. TMT SPS MS3 Mode and Close-Out features were both enabled, with a maximum of 2 peptides per protein and a maximum search time of 100 ms.

A score threshold of Xcorr ≥ 1.4 and dCn > 0.1 were used with a precursor window of 20 ppm for z = 2 and 10 ppm for z ≠ 2. The database was processed to include reversed decoys with “##” in their names as well as contaminants with “contaminant” in their names; both types of proteins were excluded from MS3 analysis. Ions up to 50 Th below or up to 5 Th above the precursor m/z were excluded from SPS, even in the case of the TMTpro isobaric tag loss (via the Isobaric Tag Loss Exclusion feature). Ten SPS ions were selected for MS3.

MS3 was performed with an MS isolation window of 1.2 Th and an MS2 isolation window of 2 Th using higher-energy CID (HCD) with a collision energy of 55%. Reporter ions were analyzed in the orbitrap at resolution = 50k with a scan range of 110–1000 Th. A custom AGC target of 300% was used with a maximum IT of 200 ms. The whole scan cycle was limited to 3 s per cycle.

Data were searched using a Comet-based in-house pipeline. C carbamidomethylation and TMT labeling of peptide N-termini and K were set as static modifications and M oxidation was used as a variable modification. For phosphoproteomic data, S/T/Y phosphorylation was also used as a variable modification and the phosphoric acid neutral loss was considered during database search. Peptide-spectrum matches (PSMs) were adjusted to a 1% false discovery rate (FDR) using a linear discriminant analysis and then assembled further to a final protein-level FDR of 1%. Peptides, proteins, and phosphorylation sites were only counted as being quantified if the sum of TMT peak signal-to-noise ratios exceeded 10 per channel; the rest were discarded. Each channel was normalized to the total sum of signal in that channel; for phosphoproteomic data, each channel was normalized to the total sum of the corresponding channel in the non-phosphoproteomic dataset.

### Proteomic and phosphoproteomic data analysis (*in vitro* CD72-treated microglia)

Significantly changing proteins and phosphosites were determined by performing Welch’s T-test between experimental and control conditions, then adjusting p-values for multiple comparisons using BH method. For input to Panther (version 19.0) GO-biological processes over-representation analysis, significantly changing proteins with adjusted p-value <0.05 and fold change >1.5 were used, while significantly changing phosphoproteins with adjusted p-value <0.05 and fold change >1.3 were used. The background was all proteins/phosphoproteins identified in the respective dataset. The overrepresentation test used Fisher’s Exact test and corrected using False Discovery Rate P<0.05. Significant pathways of interest were manually selected for visualization by bar chart.

### Molecular cloning of AAV constructs

pAAV.CAG.mNeonGreen plasmid, harboring AAV2 ITR sequences and the CAG promoter, was digested with KpnI and HindIII restriction enzymes to remove the mNeonGreen sequence and the resulting backbone was gel extracted^84^. Three fragments were prepared by PCR and gel extraction for Gibson assembly with the backbone: CD72 cDNA sequence without the stop codon from pUC19-mCD72, T2A peptide sequence, and puromycin resistance gene (both from in-house plasmids). Gibson assembly was performed using NEBuilder Hifi DNA Assembly Master Mix according to manufacturer’s protocol. For the control AAV genome plasmid, the same was performed, except using a 3xFlag tag sequence instead of CD72 cDNA sequence.

### AAV production

AAVs were generated by triple transfection of adherent HEK293T cells using polyethylenimine (PEI). Three days after transfection, cells were harvested and AAVs were purified using ultracentrifugation over iodixanol gradients, and titered as previously described^85^.

### AAV transduction of mixed glia cultures

Mixed glia derived from Cx3cr1-GFP mice were cultured from frozen in 6 well plates for 10 days, treated with AAV expressing CD72 or 3xFlag control for two days, then selected using 2µg/ml puromycin for two days.

Transduced microglia were enriched using MACS (see <u>Recombinant protein screen in mixed glia culture</u>). For western blotting analysis, microglia were collected into a cell pellet, flash frozen with liquid nitrogen, and stored at -80°C until further use. For tiling analysis, microglia were recombined with uninfected, non-microglia from a separate well at 10% microglia and replated to a 96 well plate coated with Matrigel at 50K total cells per well (5,000 microglia; 45,000 non-microglia) and returned to the incubator for 2 days. Cultures were fixed and stained according to <u>Fixation and immunofluorescent staining of recombinant protein screen</u> then imaged according to <u>Automated imaging of recombinant protein screen.</u> Image analysis was performed according to <u>Image processing and analysis of tiling features.</u>

### Western blotting analysis

Pellets were lysed in 100µl of cold 0.2% SDS RIPA buffer: 50 mM Tris-HCl [pH] 8.0, 150 mM NaCl, 0.2% SDS, 0.5% sodium deoxycholate, 1% Triton X-100, 1x protease inhibitor cocktail, 1mM PMSF. The samples were incubated for 10 minutes on ice, then DNA was sheared using a 1ml syringe with a 27-gauge needle 10 times up and down. Samples were incubated at 4°C while rotating for 30 minutes, then cleared via centrifugation at 16,000g for 10 minutes at 4°C. Supernatant was transferred to a new tube on ice. BCA assay was performed on the lysate to determine total protein concentration. 20µg of total protein was prepared in 20µl of NuPage LDS sample buffer with 40mM DTT final. Samples were heated at 70°C for 10 minutes then loaded onto 4-12% bis-tris protein gel, then transferred to PVDF membrane. The membrane was blocked with 3% BSA in PBST (0.1% Tween20) for 1 hour at room temperature. Primary antibodies: rat anti CD72 1:1000 or mouse anti β-actin 1:10,000 were incubated in 3% BSA in PBST overnight at 4°C while rocking. The membrane was washed three times with PBST for 10 minutes each while rocking, then peroxidase-conjugated secondary antibodies: donkey anti mouse IgG 1:5,000 or anti-rat IgG 1:5,000 were incubated in 3% BSA in PBST for 1 hour at room temperature while rocking. The membrane was washed again, then dipped in Clarity Western ECL substrate for one minute and imaged. Quantification of band density was performed in Fiji.

