## Supplemental Figures for "Cell-surface proteomic profiling identifies CD72 as a regulator of microglial tiling"

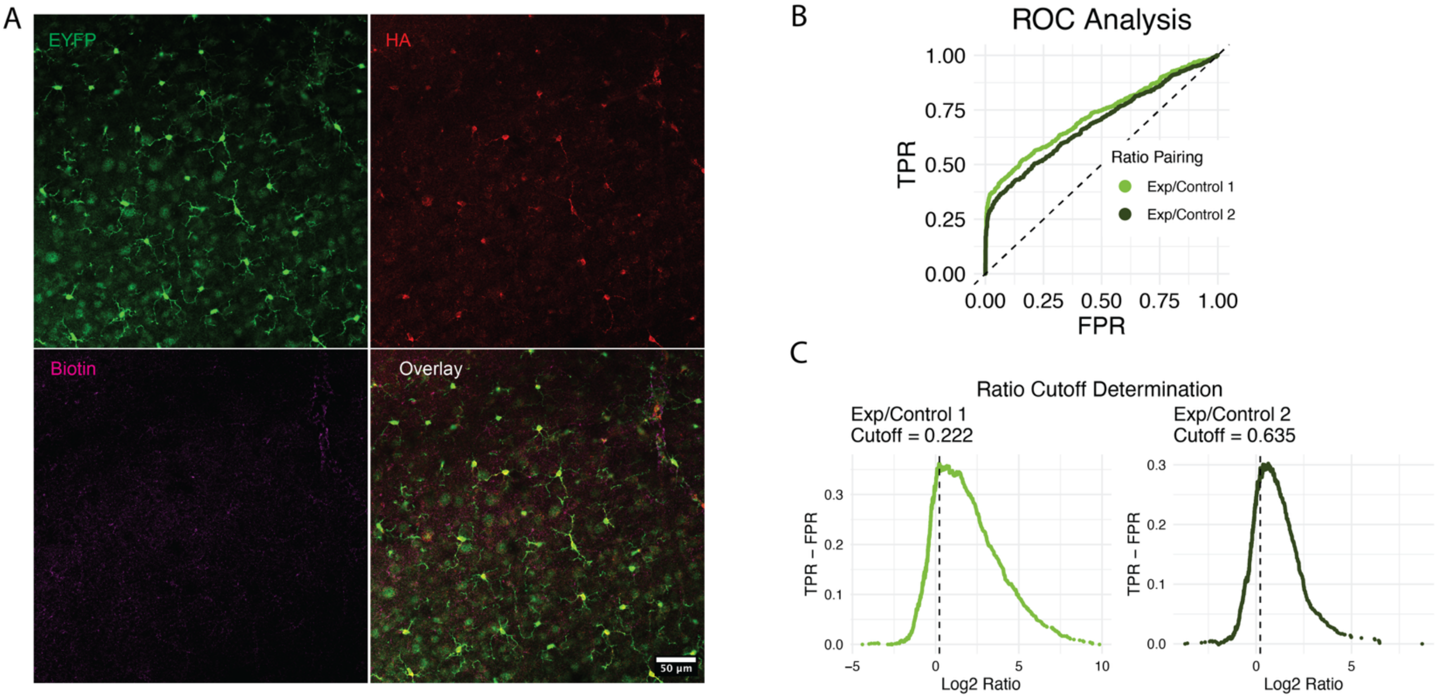


**Figure S1. Proximity labeling in acute brain slices and ratiometric cutoff determination for *in vitro* cell-surface proteome analysis**

**(A)** Confocal images of 250µm acute brain slices from Cx3cr1-Cre$^{ERT2}$-EYFP; Cre-iPEEL mice, treated with all proximity labeling reagents *in situ* and stained for microglia (EYFP, green), HRP expression (HA-tag, red), and biotinylated protein (Streptavidin-647, magenta). **(B)** Receiver operating characteristic (ROC) analysis showing true positive rate (TPR) against false positive rate (FPR) for both ratio pairings (Experimental/Control 1 and Experimental/Control 2) devised during ratiometric cutoff determination for *in vitro* cell-surface proteome analysis. **(C)** Cutoff was determined for each ratio pairing where TPR-FPR was maximal. Proteins with ratios above the cutoff were retained. See also Figure 1F, Table S1, and STAR Methods.

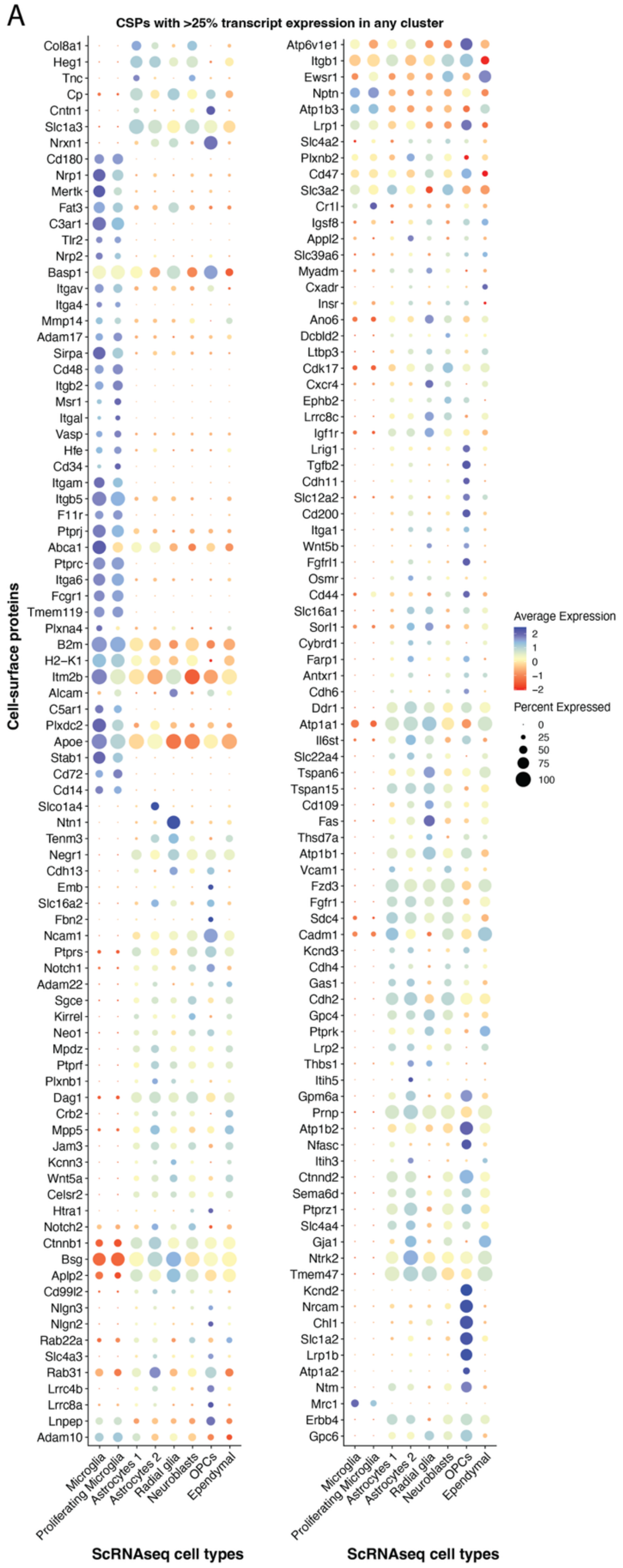


**Figure S2. Cell type origins of microglial cell-surface proteome**

**(A)** Dot plot of a subset of CSPs identified in Figure 1 (y-axis), filtered for only those with corresponding transcript expressed by >25% of cells within any cluster (identified in scRNAseq, along x-axis). Order of proteins determined by hierarchical clustering.

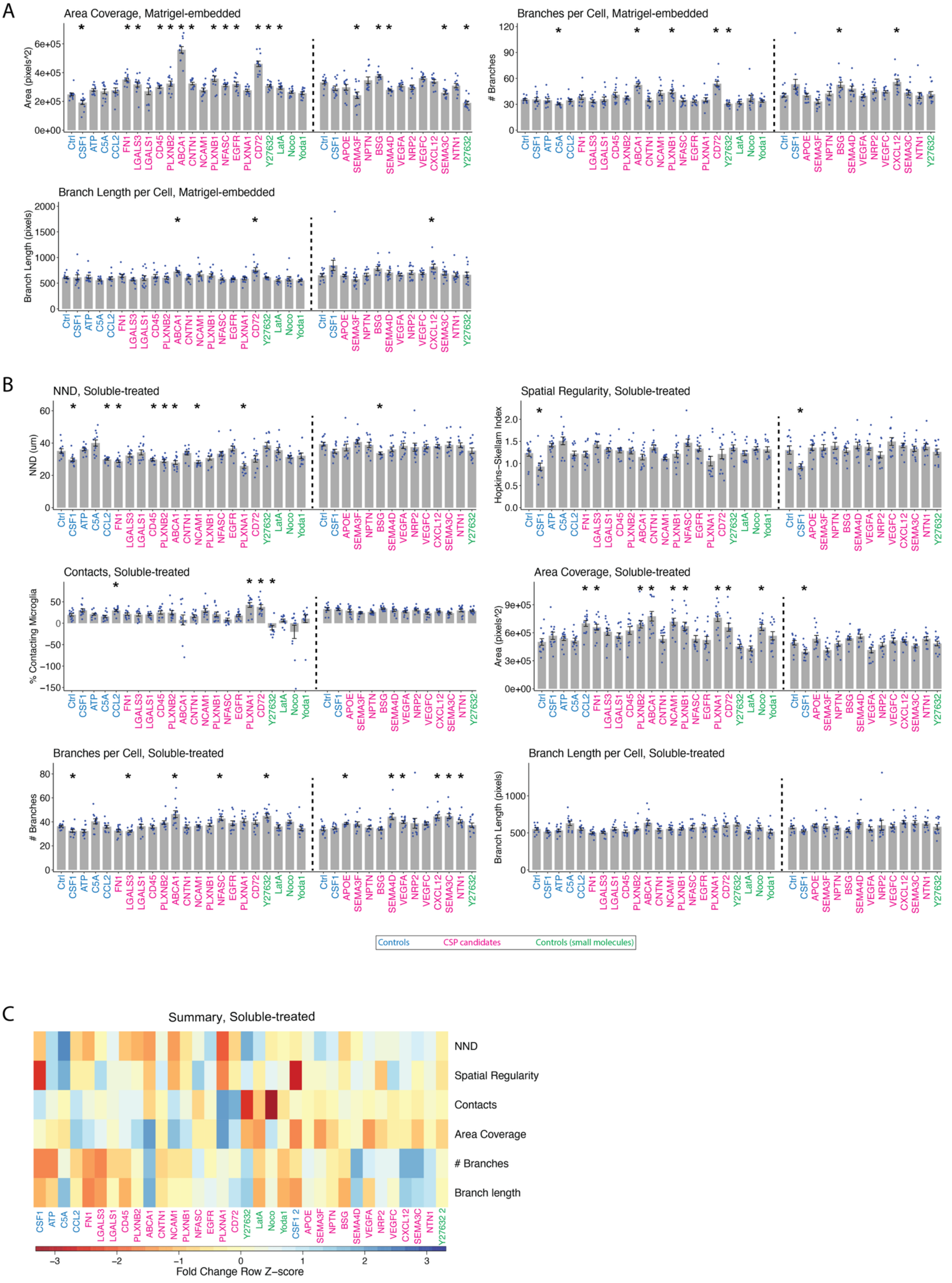
A
Area Coverage, Matrigel-embedded
Area (pixels^2)
Branches per Cell, Matrigel-embedded
### Branches
Branch Length per Cell, Matrigel-embedded
Branch Length (pixels)
B
NND, Soluble-treated
NND (um)
Spatial Regularity, Soluble-treated
Hopkins−Skellam Index
Contacts, Soluble-treated
% Contacting Microglia
Area Coverage, Soluble-treated
Area (pixels^2)
Branches per Cell, Soluble-treated
### Branches
Branch Length per Cell, Soluble-treated
Branch Length (pixels)
Ctrl CSF1 ATP C5A CCL2 FN1 LGALS3 LGALS1 CD45 PLXNB2 ABCA1 CNTN1 NCAM1 PLXNB1 NFASC EGFR PLXNA1 CD72 Y27632 LatA Noco Yoda1
Ctrl CSF1 APOE SEMA3F NPTN BSG SEMA4D VEGFA NRP2 VEGFC CXCL12 SEMA3C NTN1 Y27632
Controls
CSP candidates
Controls (small molecules)
C
Summary, Soluble-treated
NND
Spatial Regularity
Contacts
Area Coverage
### Branches
Branch length
CSF1 ATP C5A CCL2 FN1 LGALS3 LGALS1 CD45 PLXNB2 ABCA1 CNTN1 NCAM1 PLXNB1 NFASC EGFR PLXNA1 CD72 Y27632 LatA Noco Yoda1 CSF1 2 APOE SEMA3F NPTN BSG SEMA4D VEGFA NRP2 VEGFC CXCL12 SEMA3C NTN1 Y27632 2
Fold Change Row Z−score

**Figure S3. An image-based screen identifies CD72 as a strong regulator of tiling**

**(A)** Data from area coverage, branches per cell, and branch length per cell from the Matrigel-embedded screen. N= four images in each of three wells, 12 technical replicates. Bars show mean ± SEM. Two rounds of screening were performed, with rounds separated by vertical dotted lines. Molecules are shown in the order they were tested. X-axis text color represents category of molecule. Adjusted p-values (BH method): *$p<0.05$ by Welch's t-test between respective molecule and Ctrl (vehicle 0.1% BSA) within round. See also Figure 3D. **(B)** Data from all six features from the soluble-treated version of the screen. Details same as (A). **(C)** Summary heatmap of all features for every molecule in both rounds of the soluble-treated screen. Color bar represents fold change over respective Ctrl within round. X-axis text color represents the same as in (A) and (B).

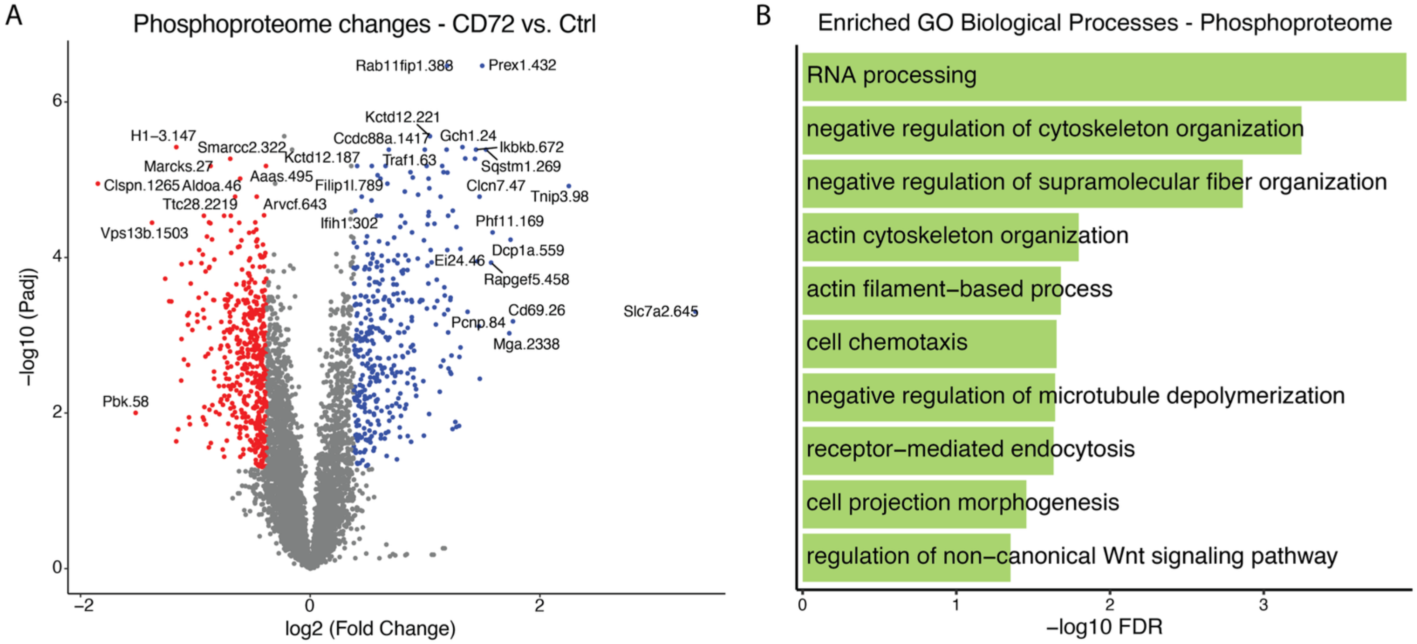


**Figure S4. rCD72 induces molecular pathways consistent with tiling disruption and microglial immune response**
**(A)** Volcano plot showing phosphosite occupancy changes in microglia enriched from the mixed glia culture treated with Ctrl (vehicle 0.1% BSA) or 400ng/ml rCD72, N=6 each. Welch's t-test was applied between Ctrl and rCD72, then p-values adjusted using BH method. Average fold changes were calculated. Red dots denote downregulation and blue dots denote upregulation with rCD72 treatment (fold-change cutoff: 1.3, adjusted p-value cutoff: 0.05). Labels formatted as "gene.site." See also Table S6. **(B)** Enriched biological processes from significantly changing phosphoproteins.
